# From 2D Monolayer to 3D Spheroid: Matrigel’s Spatial and Chemical Cues Remodel the Calu-3 Proteome

**DOI:** 10.1101/2025.08.28.672967

**Authors:** Ariel M. Maia, Luiz A. Basso, Pablo Machado, Cristiano V. Bizarro

## Abstract

The limited physiological relevance of conventional 2D cell cultures hampers translational research. While 3D cultures offer a superior alternative, their reliance on undefined scaffolds like Matrigel makes it difficult to distinguish cellular responses to physical architecture from those to biochemical cues. This study provides a high-resolution, label-free quantitative comparison of the Calu-3 lung adenocarcinoma proteome in 2D monolayers versus 3D Matrigel-embedded spheroids. To further dissect the specific contributions of 3D architecture versus biochemical cues from the matrix, we included an intermediate condition where a 2D monolayer was cultured in contact with Matrigel droplets (2M). After a stringent data analysis pipeline that controlled for matrix contamination, we quantified over 3,900 human proteins and revealed that 3D culture architecture is the dominant driver of proteomic remodeling. We identified 72 differentially enriched and 48 conditionally quantified proteins distinguishing the 3D spheroids from monolayers, functionally characterized by a down-regulation of cell-surface and cholesterol biosynthesis proteins and an up-regulation of proteins associated with iron clusters. While direct comparison between the two bidimensional cultures yielded no significant differences, a dedicated paired statistical analysis demonstrated that biochemical cues from Matrigel alone prompted a subtle but significant proteome-wide shift, moving the cells closer to the 3D phenotype. This study delivers a detailed blueprint of the profound proteomic reprogramming induced by 3D culture and, by isolating the matrix’s biochemical influence, provides a critical resource for rational model selection and the design of defined synthetic matrices.

## INTRODUCTION

The successful translation of preclinical research to clinical applications fundamentally depends on the physiological relevance of *in vitro* models. Traditional bidimensional (2D) cell cultures, while foundational to cell biology, grow as monolayers on artificially rigid substrates, a state that fails to recapitulate native tissue architecture and has been shown to favor non-epithelial cell states (1). This is a critical limitation, particularly in cancer research, the field where this culture is most advanced, as most human cancers are carcinomas derived from epithelial tissues. Consequently, there has been a shift towards three-dimensional (3D) culture systems, such as spheroids and organoids, which better preserve epithelial phenotypes, cell-cell interactions, and tissue-specific functions, thus providing more reliable models for studying disease pathogenesis and drug response (2). This move is further underscored by recent initiatives from regulatory bodies like the FDA and NIH to prioritize robust non-animal models for new drug development (3, 4).

One of the methods used to grow those advanced 3D models, however, is heavily reliant on the use of hydrogels, most commonly animal-derived extracellular matrix extracts like Matrigel, a basement membrane extract derived from mouse Engelbreth-Holm-Swarm (EHS) tumors. While Matrigel has been the gold standard for inducing self-organization and maintaining cell viability, especially in the field of stem-cell research, its use presents significant challenges. It is a biochemically undefined mixture with high batch-to-batch variability, it is costly, and its animal origin can be a hurdle for clinical translation (5, 6).

While 3D Matrigel cultures tend to be superior over 2D plastic, the individual contributions of the 3D physical architecture *versus* the biochemical and mechanical cues of the Matrigel substrate are not well quantified (7, 8). Pinpointing these effects is essential not only for the development of synthetic alternatives that can replicate the benefits of ECM without the associated drawbacks, but in the meantime for a well substantiated use of this technology in research and medical applications (6). To address this gap, we employed a quantitative proteomic approach using the Calu-3 lung adenocarcinoma cell line, a widely used model in respiratory and cancer biology. We established three distinct culture conditions: a standard 2D monolayer, a 2D monolayer cultured in contact with Matrigel-GFR droplets (2M), and 3D spheroids fully embedded in Matrigel-GFR (3D).

Here, we provide a high-resolution proteomic characterization and comparison of these three states. We demonstrate that while 3D architecture is the dominant driver of proteomic reprogramming, the biochemical cues from the Matrigel substrate alone are sufficient to induce a subtle but statistically significant shift towards the 3D phenotype. This dataset provides a critical resource for researchers, offering a detailed rationale to guide the processing and selection of appropriate culture models and contributing fundamental knowledge towards the development of defined, synthetic matrices for lung cell biology.

## EXPERIMENTAL PROCEDURES

### Experimental Design and Statistical Rationale

This study aimed to quantitatively assess proteomic alterations in Calu-3 cells exposed to distinct microenvironments: standard 2D culture (2D), 2D culture in contact with Matrigel-GFR (2M), and 3D culture fully embedded within Matrigel-GFR (3D). The objective was to differentiate proteomic signatures primarily influenced by soluble/contact-dependent Matrigel cues from those driven by immersive 3D spatial architecture.

A total of 18 samples (3 experimental conditions x 3 biological replicates x 2 technical replicates) were analyzed. n=3 is a widely accepted standard in discovery proteomics, providing sufficient statistical power to detect the changes, especially in the experimental design of this work. For statistical analysis, the intensities from the two technical replicates were summarized into a single protein intensity value for each of the 9 biological samples (3 per condition). The number of biological and technical replicates was set based on the availability of the proteomics facility to process it. The same flask of Matrigel was used in all biological and technical replicates. Samples were analyzed by LC-MS/MS in DDA mode, processed by the FragPipe LFQ-MBR workflow, possible contaminants filtered out and ion intensities summarized by directLFQ to yield a single protein intensity value per biological replicate prior to statistical analysis. All experimental conditions were cultured and harvested in parallel to minimize batch effects related to handling.

Protein intensity data derived from directLFQ were filtered to include proteins quantified in all replicates across all conditions (“non-missing proteins”) or those with at most one missing value per condition (“partially-missing proteins”) which were then imputed using the seqKNN algorithm (k=10) based on the "non-missing" protein distribution in an effort to retain more proteins for downstream analysis.

Differential protein expression between conditions (2D vs. 2M, 2D vs. 3D, and 2M vs. 3D) was assessed using Limma implemented via FragPipe-Analyst (v1.13). P-values were adjusted for multiple testing using the Benjamini-Hochberg procedure to control the false discovery rate (FDR). Proteins were considered significantly differentially enriched (DEPs) if they exhibited an adjusted p-value < 0.01 and an Log_₂_FC ≥ 1.

In addition to DEPs, Conditionally Quantified Proteins (CQPs) were identified based on consistent quantification in all replicates of one or two conditions and complete absence of quantifiable signal in all replicates of the remaining condition(s), coupled with an intensity filter (mean intensity above the 5th percentile of the “non-missing dataset”) to enhance confidence and reduce probability of the absent signal coming from being too close to the MS1 limit of detection. To test the hypothesis that the 2M condition represents an intermediate phenotype, a one-sided, paired t-test was performed on the absolute Log₂(Fold Change) values of the 2D vs 3D and 2M vs 3D comparisons.

The list of DEPs and CQPs genes were analyzed in StringDB v12.0 (9). Protein-protein interaction and network analysis was generated using a minimum required interaction score of high confidence (> 0.7). Functional enrichment for Gene Ontology (GO) terms, KEGG and WikiPathways, and UniProt Annotated Keywords was conducted using an FDR significance threshold of < 0.01 against the background list of all proteins quantified used in the differential analysis.

### Cell culture

Calu-3 cells (BCRJ 0264 - ATCC HTB-55) were acquired from Rio de Janeiro’s Cell Bank, in Brazil. Cells were maintained in a T75 flask with Dulbecco’s Modified Eagle Medium (DMEM) supplemented with 1.8 g/L NaHCO₃, 10 % (v/v) heat-inactivated fetal bovine serum (FBS), 100 U/mL penicillin, 100 µg mL⁻¹ streptomycin and 0.1% (v/v) amphotericin B (“complete medium”) at 37 °C in 5% CO₂. Medium was replaced every 2 days and cells were subcultured at ∼80% confluence (every 4 to 5 days).

On the day before the experiment, the cell’s media was changed one last time. For seeding, cells were washed twice with 5mL of Dulbecco’s Phosphate-Buffered Saline (DPBS) and detached using 3 mL of Trypsin-EDTA (10 min, 37 °C). After quenching with 7mL of complete medium and centrifugation (200 x g, 5 min, 25 °C), the pellet was resuspended with complete medium, cells were counted, split into 2 tubes (2D/2M and 3D) and centrifuged again.

A 6-well plate was used for the experiment. For the 2D and 2M groups, cells were resuspended with complete medium and plated 1 × 10⁵ cells per well in 3 mL. This initial cell seeding density was optimized to reach 100% confluency only at the experiment’s end, preventing senescence from affecting the analysis. In 2M, two 50 µL Matrigel-GFR droplets (7 mg/mL) had been pre-polymerized (30 min, 37 °C). At the same time, the 3D group pellet was resuspended in the same diluted Matrigel-GFR. Two 50 µL droplets containing 5 × 10⁴ cells each were dispensed per well and allowed to solidify (30 min, 37 °C) before covering it with 3 mL complete medium. This Matrigel concentration was chosen to enhance the reproducibility of this experimental design as Matrigel-GFR batches’ lower range concentration is 7 mg/mL. To reduce serum interference in cell’s expression and background noise in LC-MS/MS (10, 11), FBS was lowered to 5% 24h post-seeding. This concentration was previously validated for long-term Calu-3 cell culture (12). Medium was changed every 48 h, and cells were harvested on day 7.

### Cell harvest

On day 7 of the culture cells had just reached 100% confluency and were harvested. Dispase II was used to dissolve Matrigel from the samples as previously reported to be one the best alternatives to minimise matrix-derived carry-over in proteomics (13). The medium from all groups was discarded and wells were washed 3 times with 1 mL room-temperature DPBS. 2 mL of warm dispase II solution (2 mg/mL = 5 IU/mL, 0.22 µm-filtered in DPBS) was added to 3D wells, while the same volume of DPBS was added to wells 2D and 2M, and left to incubate at 37°C for 30 minutes. After that, the Matrigel domes from the 3D wells were physically disrupted with a p1000 tip by gently pipetting up and down 5 times. The Matrigel domes from 2M were carefully aspirated from the plate without disrupting the monolayer, supernatants from 2D and 2M wells were discarded and 3 mL of the dispase II solution was added in each well. The plate was put back into the incubator for another 30 minutes.

After the incubation, cells from the 2D and 2M well were gently detached using a scraper and all groups were transferred along with the supernatant into a 15 mL falcon, where 7 mL of cold DPBS was added in order to dilute the dispase II solution. Cells were then centrifuged for 10 minutes at 500 g at 4°C, the supernatant discarded, pellet was resuspended into 2 mL of cold DPBS, transferred into a 2 mL eppendorf and centrifuged again for 5 minutes at 500 g at 4°C. The pellet was washed and centrifuged 3 times in order to reduce remaining traces of Matrigel and FBS, and in the last time the supernatant was discarded and the pellet-containing eppendorfs were stored in a -80° C freezer.

### Cell lysis, protein extraction and quantification

All samples were submitted to the same protein extraction protocol from Vernavides et al (14). Briefly, cell pellet samples were homogenized with 200 µL of a lysis buffer made of 8 M urea, 50 mM Tris-HCl and 1 mM EDTA at pH 8.5 with milli-Q water. The samples were placed on ice and sonicated two times for 20 seconds, with a 40 seconds pause in-between, using a Qsonica’s “Q125 Sonicator®” equipped with a probe #4423 of 5/64” (2 mm). The sonicated samples were then centrifuged at 12.000 g for 20 minutes at 10°C, the supernatant collected into a new eppendorf and stored at-80°C. Protein concentration was determined by BCA method (15) (ThermoFisher) after diluting 30 µL of the sample solution in 50 µL of milli-q water to reduce urea concentration from 8M to 3M, preventing interference with the BCA quantification as per manufacturer reagents compatibility instructions.

### In-solution digestion and sample preparation

All samples were submitted to the same protocol of in-solution digestion as described previously (16). Briefly, 18 µg of protein from each sample was aliquoted and volume was equalized to 40 µL with 8 M urea. Proteins were reduced with 5 mM dithiothreitol (DTT) for 25 minutes at 56 °C and alkylated with 14 mM iodoacetamide (IAA) for 30 minutes at room temperature in the dark. The remaining IAA was removed by adding excess DTT. To reduce the final concentration of urea to 1 M, the mixtures were diluted with a 50 mM ammonium bicarbonate buffer. Proteins were digested with trypsin (1:50, w/w) for 18 hours at 37°C, and then 3% formic acid (v/v) was added to stop the digestion. The tryptic peptides were desalted with C18 StageTips (17) and samples were vacuum dried for 18 hours, then stored at -20°C.

### LC-MS/MS analysis

The samples were analyzed by LC-MS/MS on an Orbitrap Exploris 240 mass spectrometer (Thermo Fisher Scientific, USA) connected to the EASY-nLC system 1200 (Proxeon Biosystem, USA) through a Proxeon nanoelectrospray ion source. Peptides were separated by a 2-40% acetonitrile gradient during 155 minutes, using 80% acetonitrile in 0.1% formic acid, in a trap Acclaim PepMap 100 nanoViper 2PK C18 (2 cm × ID75 μm, 3 μm particle size, Thermo Scientific) in line with an analytical PepMap RSLC C18 ES 902 column (50 cm x id 75 μm, 2 μm particle size) at a flow rate of 250 nL/min. The nanoelectrospray voltage was set to 1.7 kV and the source temperature was 275°C. The full scan MS spectra (m/z 375-1,500) were acquired in the Orbitrap analyzer after accumulation to a target value of 3 e6. Resolution in the Orbitrap was set to r = 60,000 and the 20 most intense peptide ions with charge states ≥ 2 were sequentially isolated to a target value of 3e5 and fragmented in the high-energy collisional dissociation HCD (normalized collision energy of 27%). The signal threshold for triggering an MS/MS event was set to 1 e4. Dynamic exclusion was enabled with an exclusion duration of 20 s and repeat count of 1. Maximum injection time was 60 ms.

### Raw Data Processing and Database Search

All thermo.raw files were first assessed for quality control using RawTools (v2.0.7) (18) and then converted to an open-format (.mzML) (19) using the msConvert feature of the ProteoWizard v3.0 package (20) with the “peakPicking” (centroiding) and “zeroSamples” settings, in that order.

A reference database for searching was assembled using the reviewed Swiss-Prot proteomes from Uniprot’s release v2025_02. The built database had a merged human-mouse proteome to enable Matrigel traces in the samples to be identified and excluded manually after. The *homo sapiens* reference proteome had 42,518 protein entries (20,406 canonical and 22,112 isoforms) while the *mus musculus* had 25,633 protein entries (17,224 canonical and 8,409 isoforms). The FASTA files were supplemented with common contaminants from Hao Lab’s group library (21) and built using publicly available Galaxy’s workflow from our group.

The “human-matrigel workflow” (https://usegalaxy.eu/u/maia_a_m/w/human-matrigel-database-builder-for-fragpipe) merges the human and mouse proteomes FASTA files, appends the contaminants file downloaded from Hao Lab’s GitHub page (381 sequences), removes all human and mouse contaminants (151 and 26 sequences, respectively), adds reversed sequences as decoys and tags all sequences with the appropriate “contam” and “rev” headers in a format that is compatible with FragPipe tool.

Peptide and protein identification/quantification was done using FragPipe v23.0 in an Arch Linux OS with OpenJDK v17.0.15 and python v3.12 virtual environments. The log file with all the parameters settings is available at the “Data availability” section. The LFQ-MBR workflow was loaded and most parameters were left as default. The converted .mzml files were loaded in DDA mode and all technical duplicates were identified as being from the same biological replicate so as not to inflate the *n* in the pipeline workflow.

MSFragger (22, 23) v4.2 was used for database search. Precursor and (initial) fragment mass tolerance were set to 20 ppm, while precursor true tolerance was set to 5 ppm, capitalizing on Orbitrap Exploris 240 high resolution. Mass calibration and parameter optimization were enabled. The isotope error was set to 0/1/2, and 2 missed trypsin cleavage was allowed. The peptide length was set from 7 to 50, and the peptide mass was set to 500 to 5000 Da. Oxidation of methionine and acetylation of protein N termini were set as variable modifications. Carbamidomethylation of cysteine was set as a fixed modification. The maximum allowed variable modifications per peptide was set to 3. MSBooster (24) v1.3.9 and Percolator (25) v3.7.1 were enabled to perform PSM rescoring with spectral and retention time similarities based on DIA-NN model (26) v1.8.2 beta 8. Philosopher (27) v5.1.1 with PeptideProphet (28) and ProteinProphet (29) was used to estimate the identification of FDR. The PSM/peptide/ion/proteins were all filtered at 1% FDR. Quantification and MBR was performed with IonQuant (30) v1.11.9. The minimum number of ions parameter required for quantifying a protein was set to 2 (default) and intensity normalization was turned off to make it easier to identify Matrigel proteins since the similarities between conditions were already high and downstream analysis already had a normalization step. To try and restrict MBR matching to within-condition runs, the maximum number of runs used for transfer was set to 5 and minimum required correlation between the donor and acceptor run was set to 0.2. Ion-, peptide-, and protein-level MBR FDR thresholds were all set to 1%. Default values were used for all the remaining parameters.

### Matrigel contaminants filtering

To ensure the highest confidence in our human proteome quantification and to rigorously control for residual Matrigel-derived contaminants, we implemented a robust two-level filtering strategy. As mentioned above, at the experimental level, we first selected dispase for Matrigel dissolution, a method previously shown to be superior for its high efficiency in cell recovery and matrix removal, leaving minimal contaminants (13). In addition to the dispase method we also employed repeated washes of the samples with ice-cold DPBS solution to facilitate Matrigel liquefaction and subsequent removal.

The second filter was performed at the computational level. Before transforming the raw ion intensities into protein intensities we filtered possible Matrigel contaminants from FragPipe’s output combined_ion.tsv file. Our pipeline was as follows: (i) reverse-sequence decoys were removed by excluding ions by their “rev” tag; (ii) contaminants were discarded based on their “contam” tag; (iii) as a conservative initial filter, all ions with a primary assignment to *Mus musculus* in the “Protein” column were excluded; (iv) finally, to remove the most persistent and abundant known Matrigel proteins with high confidence, we filtered out any remaining ions corresponding to the 312 high-confidence Matrigel contaminant (hc-MC) proteins reported by Wang et al. 2022 (13). This final dataset, consisting of high-confidence human ions, was then used for directLFQ quantification. The script used to do this is available at the “Data availability” section.

### directLFQ intensity generation

Protein intensities for downstream analysis were computed using the directLFQ approach (31). The matrigel-filtered combined_ion.tsv file was used as *input* for the directLFQ GUI tool (v0.3.2) ran in a python 3.9 virtual environment with minimum number of ion intensities to derive a protein intensity set at 2 and default settings for all remaining parameters. The log and parameter files of the run are also available at the “Data availability” section. The resulting directLFQ intensities were merged into the *combined_protein.tsv* file from FragPipe’s output. This manipulation was done using a python v3.13 virtual environment via VS Code v.1.100.2-1 . The script used is also available at the “Data availability” section.

### Bioinformatic and statistical analysis

For differential expression analysis, protein intensities resulting from directLFQ were filtered based on data completeness across the experimental conditions (2D, 2M, and 3D) and biological replicates (3 per condition) prior to downstream analysis inspired in the criteria used on Prostar tool (32), with modifications.

Proteins with quantified intensities in all samples across all three conditions were retained (“non-missing proteins”). We also retained proteins that were quantified in at least 2 out of 3 biological replicates at most once in each of the three conditions (“partially missing proteins”). In the most permissive scenario within this filter, a protein could have one missing value in each of the 2D, 2M, and 3D conditions, representing a global missing ratio of ⅓). Missing values in the “partially missing proteins” dataset were imputed sequentially using the seqKNN algorithm (k = 10) (33), as suggested by previous research (34). This imputation relied solely on the “non-missing proteins” dataset as reference. The resulting dataset, and the scripts used, are at the “Data availability” section.

The whole, non-filtered merged dataset was fed into the R shiny web application FragPipe-Analyst v1.13 (35, 36) for differential expression analysis using Limma (37). *P* values were corrected using the Benjamini–Hochberg (BH) procedure (38) and proteins were considered significantly regulated in pairwise comparison if their adjusted *p* value was < 0.01 and their |Log_₂_ Fold-Change| exceeded ≥ 1.

To enrich this analysis, the list of differentially enriched proteins (DEP) from FragPipe-Analyst was further supplemented with proteins we identified as exhibiting condition-restricted quantification (39, 40). Conditionally-Quantified Proteins (CQPs) were operationally defined as proteins that had no quantifiable intensity across all biological replicates of one (or two) experimental condition(s), while conversely exhibiting consistently quantifiable intensities across all biological replicates of the remaining two (or one) condition(s). To minimize the possibility of that pattern arising a false-positive hit coming from MS1 detection limits, we implemented a filtering step to retain only proteins whose mean intensity from the quantified conditions was above the 5th percentile of all protein intensities that had no missing values across all samples (“non-missing dataset”). This cutoff was chosen because, in our specific context, it represented an inflection point in the protein intensity rank plot and effectively separates the bulk of the consistently quantified proteins from those with high rates of missingness. Proteins identified as CQPs were therefore required to have an average Log₂ intensity above this threshold in the condition where they were quantified. This ensures that CQPs represent proteins that are not only consistently detected but also detected at a meaningful ion intensity level, as any protein quantified below this threshold would, in any case, be considered significantly down-regulated relative to the proteome average.

Hierarchical clustering analysis of fold-changes of the differentially enriched proteins was performed using Euclidean distance and z-score. Since the filtering criteria of the conditionally quantified proteins relied on a categorical binary presence/absence across conditions, hierarchical clustering of those proteins was done using the Jaccard method.

The list of DEPs and CQPs genes were analyzed in StringDB v12.0 (9). Protein-protein interaction and network analysis was generated using a minimum required interaction score of high confidence (> 0.7). Functional enrichment for Gene Ontology (GO) terms, KEGG and WikiPathways, and UniProt Annotated Keywords was conducted using an FDR significance threshold of < 0.01 against the background list of all proteins quantified used in the differential analysis.

To test the specific hypothesis that the 2M condition represents an intermediate proteomic state quantitatively closer to the 3D phenotype than the 2D condition, a dedicated statistical analysis was performed. This involved comparing the proteomic changes relative to the 3D condition from both monolayer perspectives (2D vs. 3D and 2M vs. 3D). First, the absolute Log_₂_(Fold Change) values from the 2D vs. 3D and 2M vs. 3D pairwise comparisons were subjected to a one-sided, paired t-test. The null hypothesis was that the mean difference between the paired absolute fold changes was zero, while the alternative hypothesis was that the changes in the 2D vs. 3D comparison were greater than those in the 2M vs. 3D comparison. To further ensure robustness against any deviations from normality, a non-parametric Wilcoxon signed-rank test was also performed on the same pairs. The practical significance of the observed mean difference was quantified by calculating the effect size using the paired Cohen’s d.

As an orthogonal approach to validate these findings, the signed Log₂(Fold Change) values were modeled using a linear regression of Log_₂_FC (2M vs. 3D) against Log₂FC (2D vs. 3D). This allowed for the assessment of the overall correlation and the determination of whether the magnitude of regulation was systematically compressed in the 2M condition. This complete battery of tests was first applied to the set of proteins identified as significantly differentially enriched and to the entire dataset of proteins that had no missing values across all samples to confirm the global trend of the results without the artifact of imputation. For these analyses, a p-value < 0.01 was considered statistically significant.

## RESULTS

### Quantitative Proteomic Profiling of Calu-3 Cells Reveals Distinct Signatures between Monolayers and Matrigel-embedded Spheroids

Using Calu-3 epithelial cells as an *in vitro* lung model, we asked how the three-dimensional micro-environment influences the cellular proteome. To disentangle these cues, cells were cultured for seven days as classic monolayers on plastic (2D), monolayers in contact with Matrigel droplets (2M) or spheroids fully embedded in Matrigel (3D). Phase-contrast images confirmed typical epithelial morphology for 2D/2M cultures and uniform, lumen-forming spheroids in 3D (Fig. 1A)

**Fig. 1.**
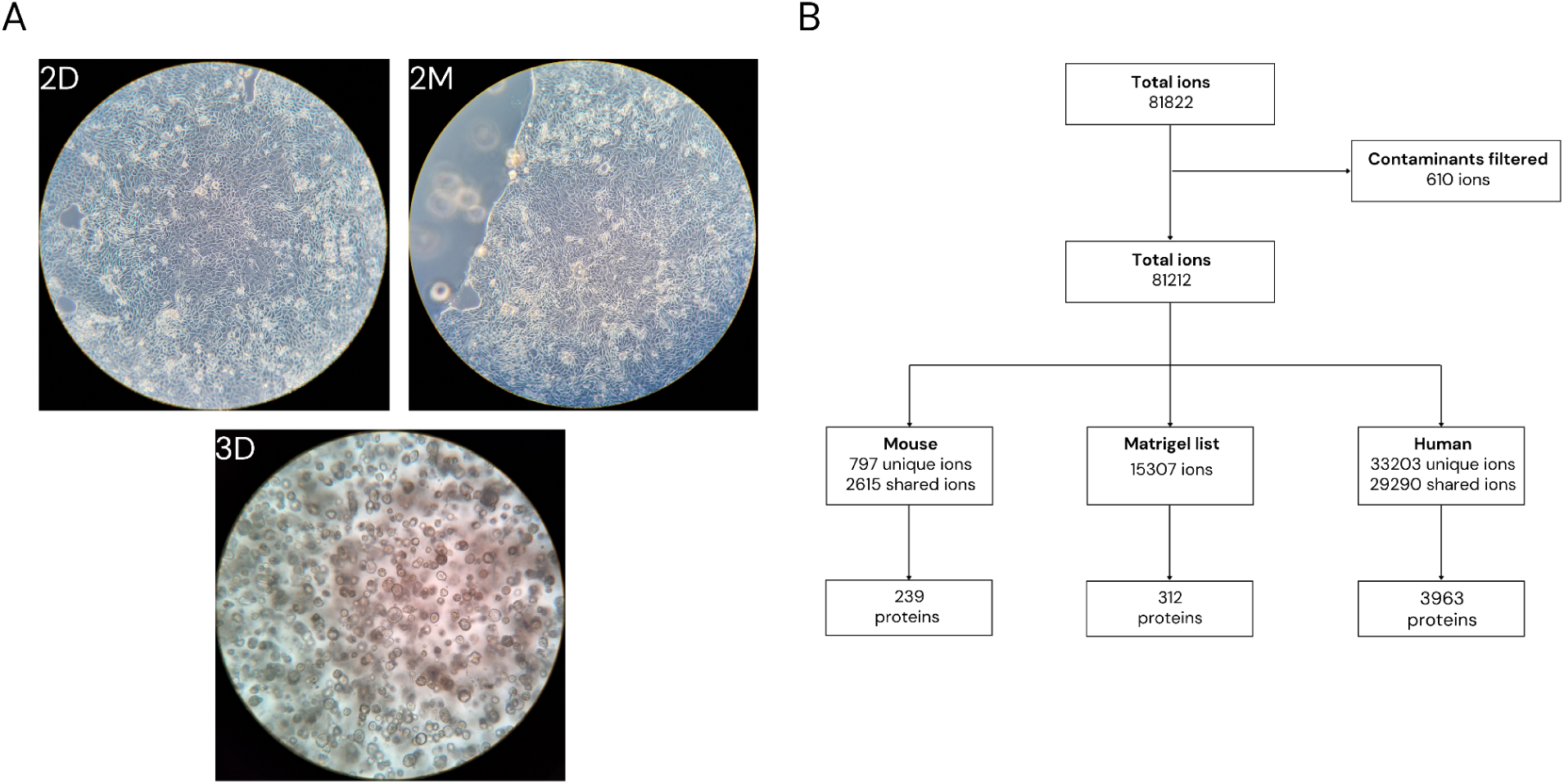
Overview of cultured conditions and Matrigel ion filtering workflow. **(A)** Representative phase-contrast microscopy images of Calu-3 cells cultured for 7 days in three distinct conditions: standard 2D monolayer on plastic (2D - top left), 2D monolayer in contact with Matrigel-GFR droplets (2M - top right), and 3D spheroids embedded within Matrigel-GFR (3D - bottom). **(B)** Flowchart illustrating the sequential filtering of ions identified by the FragPipe LFQ-MBR workflow. Starting from 81,822 non-decoy ions, common contaminants, mouse-derived ions (both unique and shared), and high-confidence Matrigel contaminants (hc-MC) were removed, resulting in a final set of 62,493 curated human-specific ions used for directLFQ protein quantification.

The FragPipe LFQ-MBR workflow search yielded 81822 ions after decoys were excluded. In an effort to reduce Matrigel interference in the analysis, a step-wise curation then discarded 610 common contaminants, 3412 mouse-derived ions (797 mouse-unique + 2615 mouse-shared) and 15307 entries in the high-confidence Matrigel panel of Wang et al. (13), leaving 62493 human ions (33203 human-unique, 29290 human-shared) for quantification (Fig. 1B).

Quantification and grouping of the curated ion list with directLFQ resulted in 3963 quantifiable proteins with at least 2 ions. Of those, 3477 proteins (87.7%) were present at least once in all conditions (Fig. 2A) and 2930 were quantified in all samples across conditions, hereafter termed “non-missing proteins” (73.9%) (supplemental Fig. S1A). This high degree of similarity and reproducibility can also be seen across biological replicates within each condition (Fig. 2B) as well as in their distribution of Log₂ intensities (supplemental Fig. S2). Missingness was also evenly distributed across 2D, 2M and 3D cultures (supplemental Fig. S3).

**Fig. 2.**
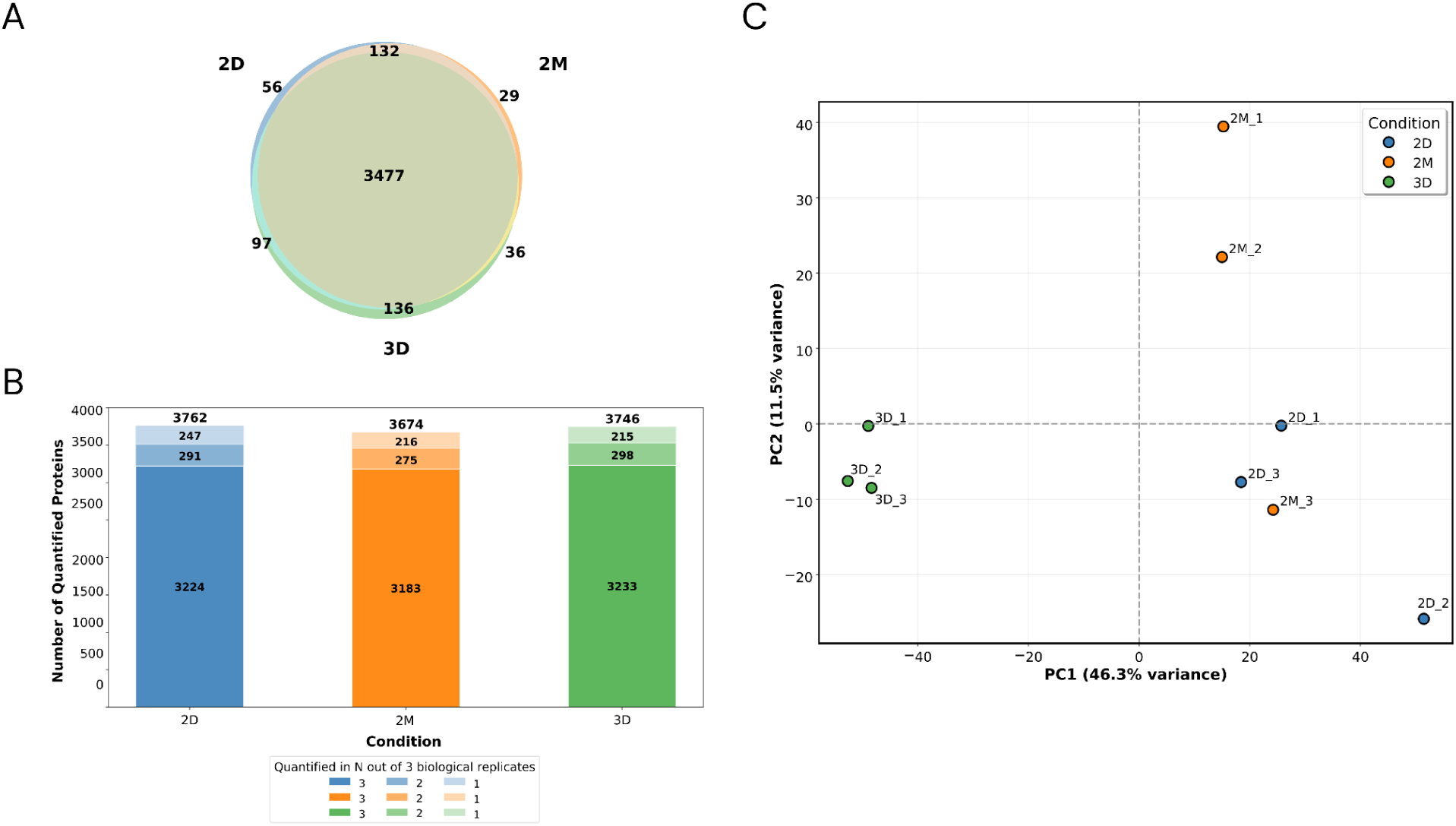
Quantitative proteomic workflow and quality assessment of Calu-3 cells cultured in 2D and 3D microenvironments. **(A)** Venn diagram illustrating the overlap of the 3,963 total proteins quantified by LC-MS/MS across the three culture conditions. A core proteome of 3,477 proteins was identified in at least one replicate of all conditions. **(B)** Stacked bar chart showing the high reproducibility of protein quantification within each condition. Bars represent the number of proteins consistently quantified in all three (dark shade), two of three (medium shade), or one of three (light shade) biological replicates (n=3). **(C)** Principal Component Analysis (PCA) of the 2930 non-missing proteins used for differential analysis across all nine samples (n=3 biological replicates per condition). PC1 explains 46.3% of the variance and clearly separates the 3D spheroids from the 2D and 2M monolayer cultures, which cluster together. PC2 explains 11.5% of the variance.

To visualize the overall proteomic relationships between the samples, a Principal Component Analysis (PCA) was performed on the 2,930 non-missing proteins (Fig. 2C and supplemental Fig. S4). The analysis using the first two components showed that culture dimensionality was the primary driver of proteomic variance, with the 3D spheroid samples forming a distinct cluster that is clearly separated from both 2D and 2M monolayer samples along PC1 (46.3% of variance). One interesting finding is that the 3D group was more tightly clustered than the bidimensional groups, suggesting less variability between biological replicates.

In contrast, clean separation between 2D and 2M groups was not observed. Although the 2M samples show a trend towards the 3D cluster along PC1, there is clear overlap, with one 2M replicate (2M_3) clustering with the 2D samples. This visual evidence suggests the proteomic effect induced by Matrigel contact alone is subtle and can be smaller than the inherent biological variability between individual monolayer replicates. Therefore, to rigorously test the hypothesis Matrigel’s biochemical components influence in the intermediate 2M phenotype, a more sensitive statistical approach beyond visual clustering was required.

### Differential Protein Expression Analysis Highlights Culture Dimensionality as Primary Driver of Proteomic Remodeling

The 2930 non-missing proteins along with 293 partially-missing proteins (at most one missing value per condition) together formed a 3223 intensity matrix. Missing values in the partially-missing subset were imputed with the seqKNN algorithm. This merged matrix was used for all Limma tests using the FragPipe-Analyst online tool. FDR correction was done using the Benjamini Hochberg method (Supplemental table S1). A Log₂FC cutoff of 1 and adjusted p-value of <0.01 was set to establish differentially enriched proteins. Using those cutoffs, we found 72 differentially enriched proteins across all conditions (Supplemental table S2). There was no significantly regulated correlation between the 2Dvs2M comparison, 68 in the 2Dvs3D comparison and 38 in the 2Mvs3D comparison (Fig. 3A). Of those, 34 were significantly quantified in both 2D/2M vs 3D comparisons, while 34 were unique only to 2Dvs3D and 4 were unique only to 2Mvs3D (Fig. 3B). Regardless of whether analyzing 2D or 2M unique proteins, the Log₂FC trends aligned with those observed in the 3D group. Interestingly, of the 34 DEPs unique to the 2D vs 3D paired analysis, 26 proteins almost met the Log₂FC >1 threshold, while passing the adjusted p-value <0.01 cutoff in the 2M vs 3D comparison.

**Fig. 3.**
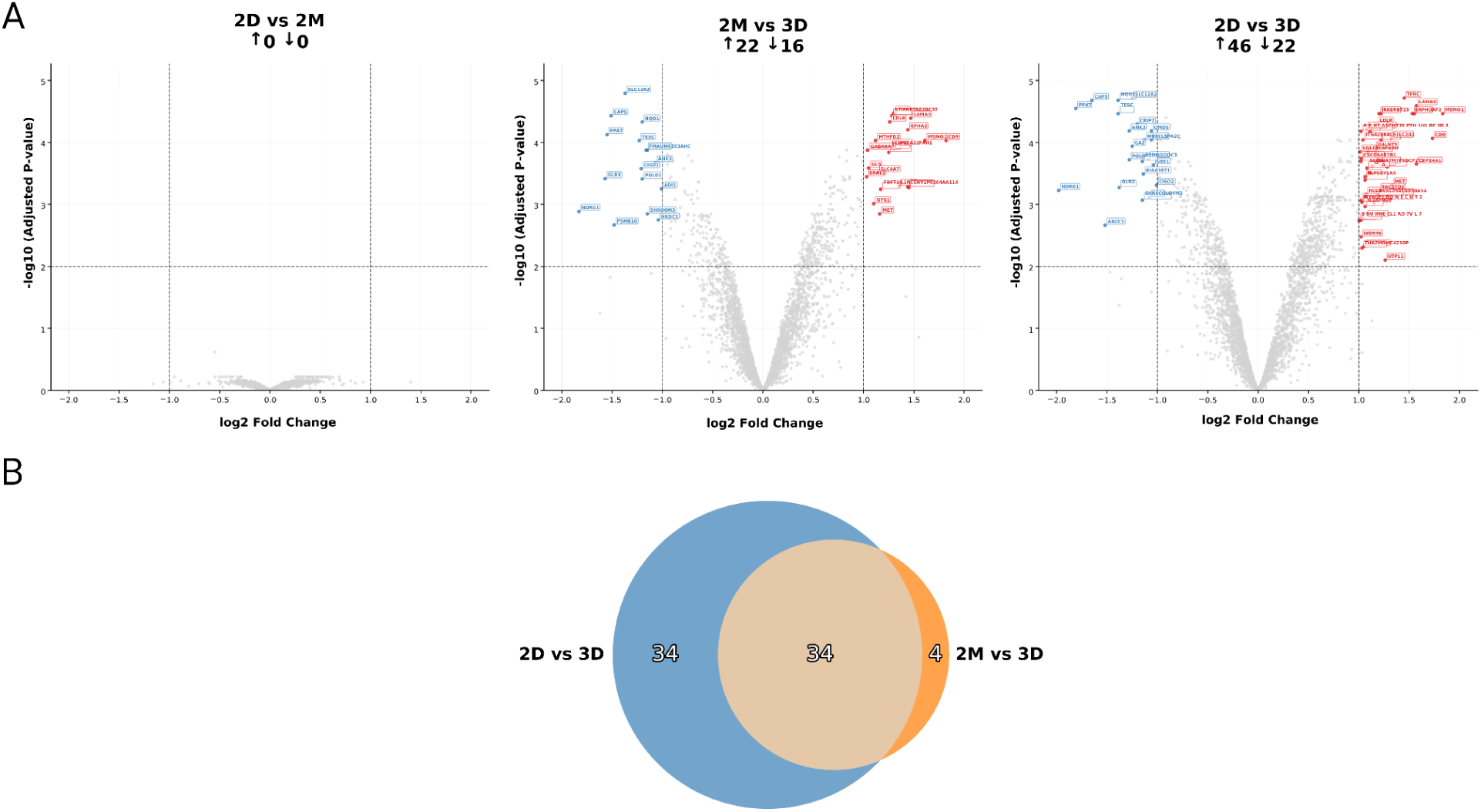
Comparative and differential protein expression between 2D, 2M and 3D conditions. **(A)** Volcano plots displaying differentially enriched proteins (DEPs) from pairwise comparisons: 2D vs 2M, 2D vs. 3D and 2M vs. 3D. The x-axis represents the Log₂(Fold Change) and the y-axis represents the -Log₁₀(adjusted p-value). Dashed lines indicate the significance thresholds (adjusted p-value < 0.01 and |Log₂FC| ≥ 1). Red and blue dots denote significantly up- and down-regulated proteins, respectively. No significant DEPs were found in the 2D vs. 2M comparison. **(B)** Venn diagram showing the overlap of the 72 total significant DEPs identified in the 2D vs. 3D and 2M vs. 3D comparisons. A core set of 34 proteins was commonly regulated in both comparisons against the 3D condition.

To further assess this we performed a euclidean hierarchical clustering with those 72 DEPs, which showed a clear clustering towards separating 3D and the bidimensional groups, with 25 proteins being up-regulated in the 3D groups (3D_up) and 47 down-regulated (3D_down) (Fig. 4A). Interestingly, the clusterization also could not separate the 2D from the 2M groups.

**Fig. 4.**
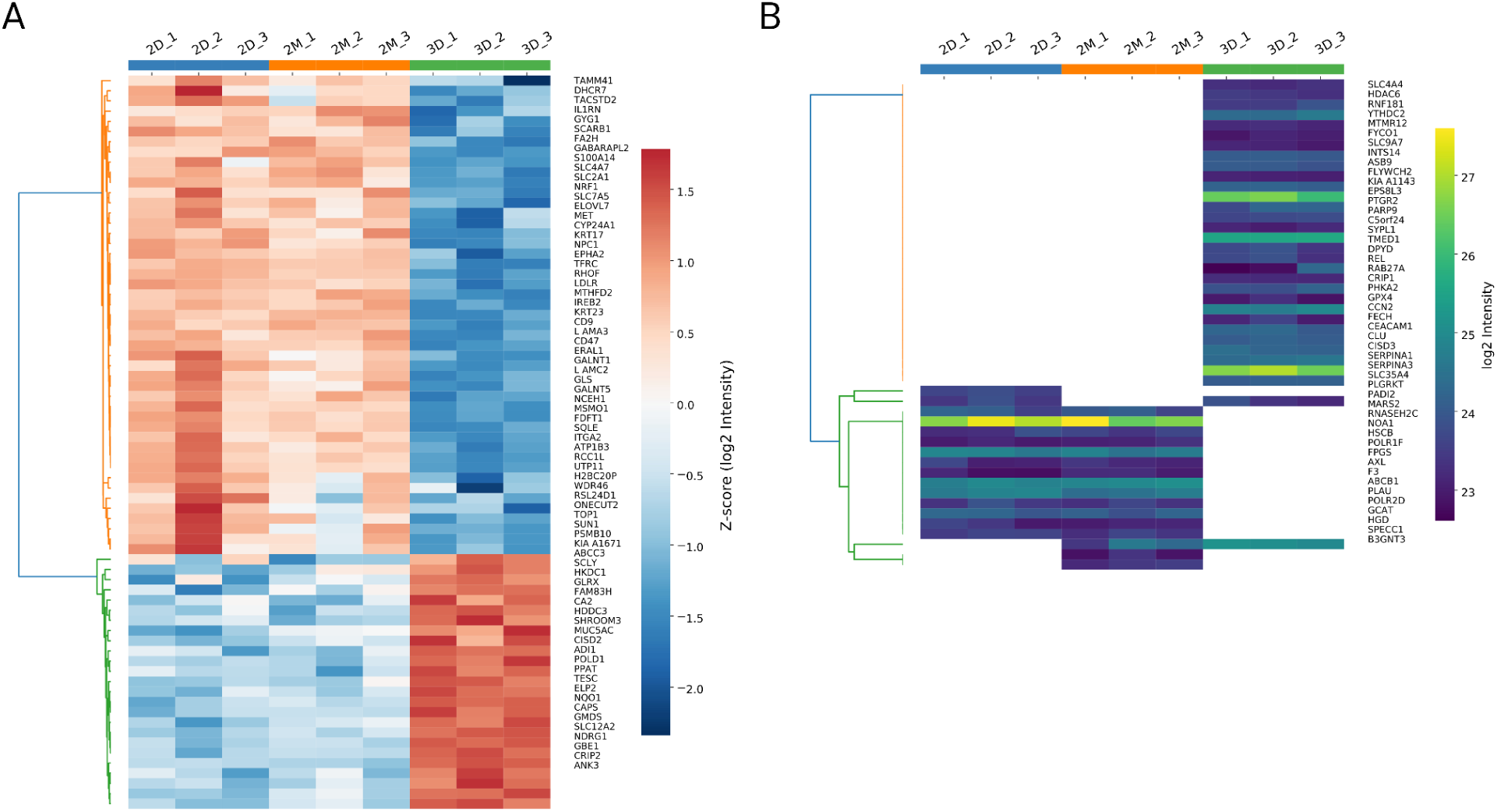
Identification and clustering of Differentially Enriched Proteins (DEP) and Conditionally Quantified Proteins (CQPs) further distinguishes 3D spheroids. **(A)** Heatmap and unsupervised hierarchical clustering of the 72 DEPs based on Z-scored Log₂(Intensity) values for each sample. The analysis reveals two primary clusters: proteins down-regulated in 3D cultures (orange cluster, n=47) and proteins up-regulated in 3D cultures (green cluster, n=25). **(B)** Heatmap showing the 48 Conditionally Quantified Proteins (CQPs) that passed an intensity filter (mean Log₂ intensity > 23.02, the 5th percentile of the non-missing dataset). CQPs are proteins consistently quantified in all replicates of one or two conditions but entirely absent in the other(s). Hierarchical clustering using the Jaccard method based on presence/absence reveals a dominant cluster of 30 proteins exclusively quantified in 3D spheroids (3D-unique) and a second major cluster of 13 proteins found only in 2D and 2M monolayers (2D/2M-unique).

### Identification and clustering of Conditionally-Quantified Proteins

To further enrich the functional analysis of the differentially enriched proteins, we sought to identify proteins that were consistently quantified in one condition but not another. We termed these “Conditionally-Quantified Proteins” (CQPs). It is important to note that the “absence” of a protein in a given condition is a technical definition, signifying that its abundance was below our stringent limit of detection, rather than a definitive claim of complete biological absence.

To identify high-confidence CQPs, we filtered for proteins that were exclusively quantified in one/two conditions while not quantified in the other two/one and passed an intensity filter corresponding to the 5th percentile of the Log₂-transformed intensity distribution of all non-missing proteins in the dataset. We found 76 proteins that fit the uniqueness presence criteria, of which 48 were retained as high-confidence CQPs after applying an intensity filter (Log₂ intensity > 23.02) (Supplemental table S3). This threshold, corresponding to the 5th percentile of non-missing proteins, was chosen to exclude low-abundance proteins prone to stochastic detection (supplemental Fig. S5A).

To identify patterns inside this new protein set, we performed a clusterization based on categorical presence/absence using the jaccard method. Similarly to the DEPs clustering, the CQP set also favored a tridimensional vs bidimensional clusterization for 43 proteins, where 30 proteins were present only in the 3D group and 13 were present only on 2D and 2M conditions (Fig. 4B). Interestingly, 5 proteins did not fit those clusters: 3 proteins showed a pattern reliant uniquely on the 2M condition and 2 based uniquely on the 2D condition.

### Functional Characterization of Significantly Regulated Proteins Altered by 3D Culture Architecture

To understand the biological implications of proteomic changes induced by the tridimensional culture environment, functional enrichment analysis was performed on two distinct gene sets using only the 3D condition as reference: the “3d_up” and “3d_down” groups. The “3d_up” group comprised 55 proteins, including 25 proteins up-regulated in the 3D group from the DEP dataset and 30 proteins conditionally quantified in the 3D group from the CQP dataset. Conversely, the “3d_down” group consisted of 60 proteins, including 47 proteins down-regulated in the 3D group from the DEP dataset and 13 proteins conditionally present in the 2D and 2M groups concomitantly. StringDB was utilized for this analysis.

For the “3d_down” gene set, 59 out of 60 proteins were found in the StringDB database. Network analysis revealed 6 distinct clusters using a high confidence minimum required interaction score of 0.7 (Fig. 5A).

**Fig. 5.**
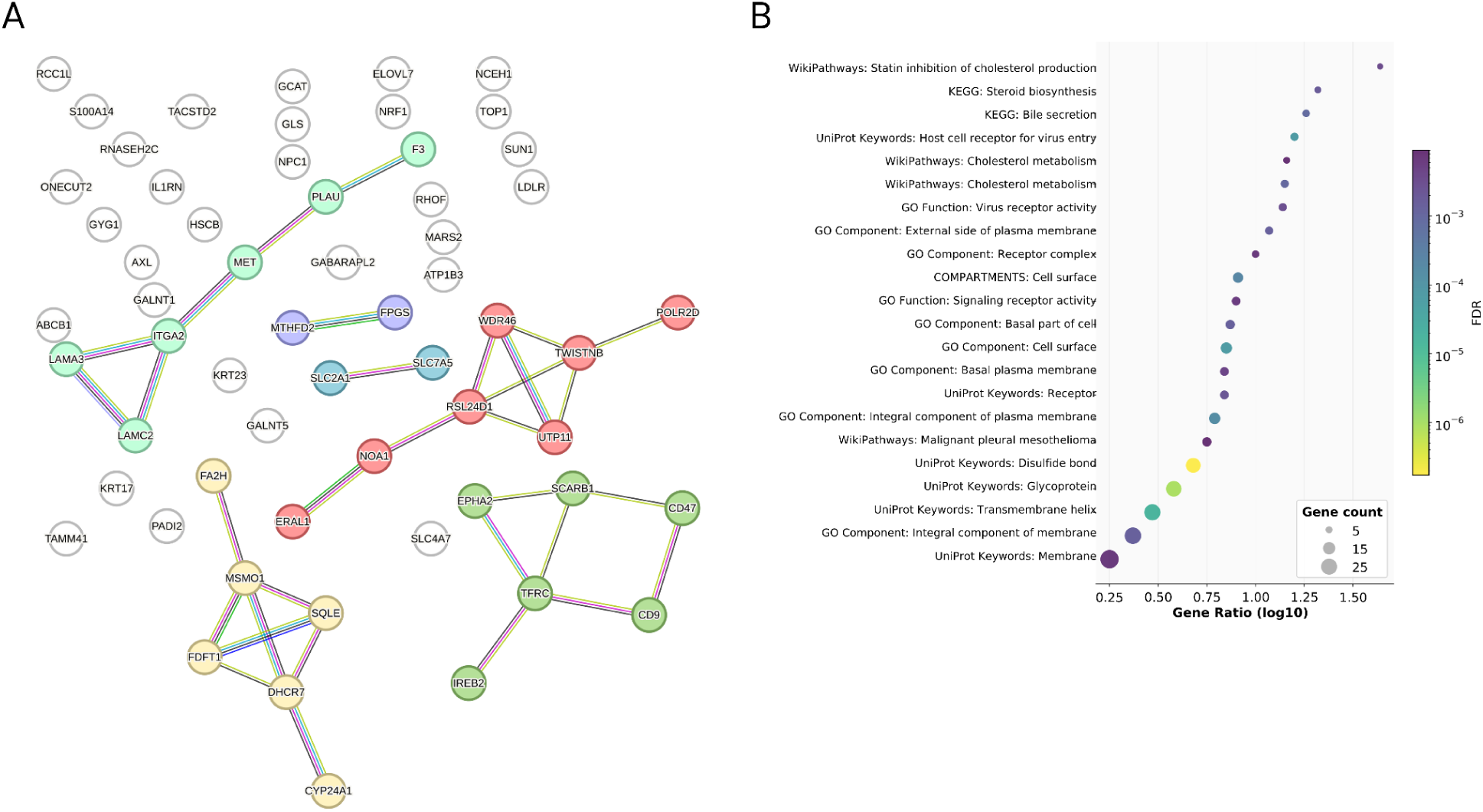
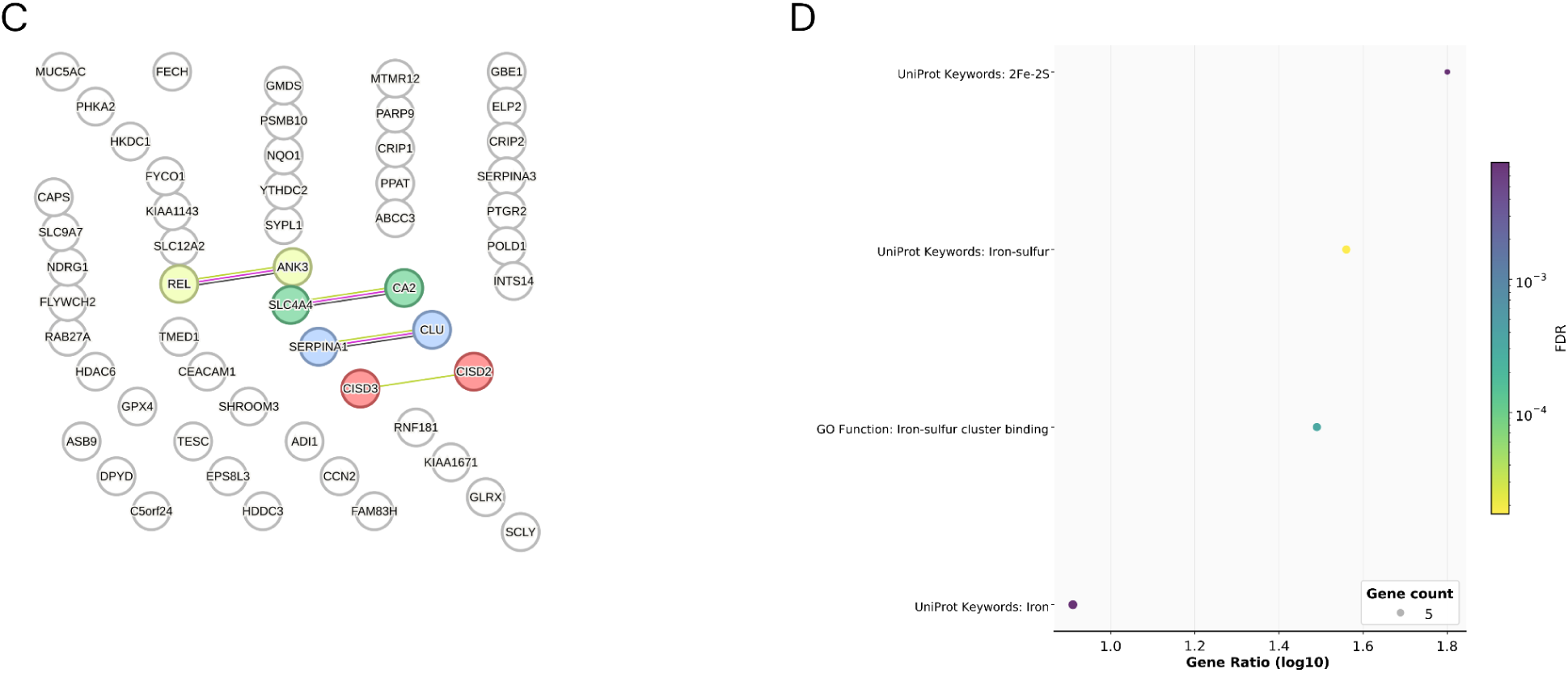
Functional enrichment analysis reveals distinct biological processes modulated by 3D culture. Protein-protein interaction networks and functional enrichment analysis were performed on proteins significantly regulated in 3D spheroids compared to 2D/2M monolayers. Networks were generated using STRING-DB (v12.0) with a high confidence interaction score (>0.7), and functional enrichment was performed for GO, KEGG, WikiPathways, and UniProt Keywords (FDR < 0.01). **(A)** Protein-protein interaction network of the 60 proteins significantly down-regulated or absent in 3D spheroids, revealing multiple interconnected clusters. **(B)** Functional enrichment analysis of the down-regulated protein set, which is significantly enriched for terms related to the cell surface, plasma membrane, signaling receptor activity, and cholesterol/steroid biosynthesis. **(C)** Protein-protein interaction network of the 55 proteins up-regulated or conditionally quantified only in 3D spheroids. The network shows fewer interactions, consistent with the inclusion of many CQPs. **(D)** Functional enrichment analysis of the up-regulated set, which is significantly enriched for iron and iron-sulfur pathways, suggesting a remodeling of metabolic or redox-related processes.

Functional enrichment analysis highlighted several significantly enriched terms (Fig. 5B). Among Molecular Functions (Gene Ontology), “Virus receptor activity” (GO:0001618, FDR = 0.0029) and “Signaling receptor activity” (GO:0038023, FDR = 0.0061) were prominent. For Cellular Components (Gene Ontology), highly significant enrichment was observed for “Cell surface” (GO:0009986, FDR = 6.73e-05), “Integral component of plasma membrane” (GO:0005887, FDR = 0.00015), and “External side of plasma membrane” (GO:0009897, FDR = 0.0014).

KEGG Pathway analysis revealed enrichment for “Bile secretion” (hsa04976, FDR = 0.00085) and “Steroid biosynthesis” (hsa00100, FDR = 0.0018). WikiPathways enrichments included “Cholesterol metabolism” (WP5304, FDR = 0.00098) and “Statin inhibition of cholesterol production” (WP430, FDR = 0.0035).

Analysis of Annotated Keywords (UniProt) showed strong enrichment for “Host cell receptor for virus entry” (KW-1183, FDR = 7.02e-05), “Disulfide bond” (KW-1015, FDR = 1.71e-07), “Glycoprotein” (KW-0325, FDR = 9.28e-07), and “Transmembrane helix” (KW-1133, FDR = 1.67e-05) For the “3d_up” gene set, network analysis in StringDB identified only 4 clusters comprising only pairs of genes, which was expected as more than half of this set came from the CQP list of genes (Fig. 5C). Functional enrichment analysis for the “3d_up” group (Fig. 5D) indicated enrichment for the Annotated Keyword (UniProt) “Iron-sulfur” (KW-0411, FDR = 0.0033), with 6 genes in the network associated with this term (CISD2, CISD3, DPYD, FECH, POLD1 and PPAT).

This detailed functional analysis indicates that the transition from 2D/2M monolayer to 3D spheroid culture induces significant remodeling of cellular processes. Down-regulation in 3D appears to prominently affect cell surface interactions, particularly those involved in receptor activity and viral interactions, alongside cholesterol/steroid biosynthesis. Conversely, proteins up-regulated or unique to the 3D environment show an enrichment related to iron cluster proteins, suggesting alterations in metabolic or redox processes requiring these cofactors.

While the global proteomic profiles showed no significant distinction between the 2D and 2M cultures in the DEPs set, a small subset of three proteins displayed expression patterns that seemed to rely on the 2M group, hinting at underlying functional differences. The proteins SPECC1 and B3GNT3 were detected in both standard 2D monolayers and 3D Matrigel cultures but were not quantified in the 2M condition. In contrast, the plasminogen receptor PLGRKT was quantified exclusively in the 2M condition. Notably, Homogentisate 1,2-dioxygenase (HGD) was quantified only in the presence of Matrigel (2M and 3D conditions), being unmeasured in the standard 2D culture. Conversely, the solute carrier SLC35A4 exhibited the opposite trend: it was identified exclusively in the 2D monolayer and was not quantified in either the 2M interface or the 3D scaffold cultures.

### Quantifying the Independent Contribution of the Matrigel Substrate to the Proteome

While Limma’s pairwise comparison between the 2D and 2M conditions did not show any significant difference in protein levels that passed the cutoffs, the PCA revealed a subtle trend that was partially obscured by biological variability, so we wondered whether the distinct conditionally-dependent examples reflected a broader but subtle systemic shift since it is known that Matrigel, even the GFR version used here, contains a myriad of growth factors and ill defined components.

To test whether the 2M condition lies in an intermediate proteomic state, we performed a paired analysis comparing the magnitude of proteomic changes of both monolayer conditions relative to the 3D state. We compared the absolute log₂ fold-changes (∣Log_₂_FC∣) between 2D vs 3D and 2M vs 3D for each of the 72 DEPs as well as for all the 2930 that were quantified in all three conditions. A paired analysis of the 72 differentially enriched proteins revealed that the absolute magnitude of expression change was on average smaller in the 2M condition compared to the 2D condition (mean |Log_₂_FC| = 1.073 vs 1.208, respectively; paired t-test, p < 0.01) (Fig. 6A). The same trend was also seen when extending the analysis to all the non-missing proteins dataset. On average, cells in 2D diverged from 3D by ∣Log₂FC∣=0.30, whereas 2M diverged by only 0.26 (Δ=0.039 log_₂_ units; paired t-test, one-sided p < 0.01; Wilcoxon signed-rank, one-sided p=1.8×10−67) (Fig. 6B). This small but consistent 0.04 log₂ unit “compression” corresponds to a 13 % smaller average fold-change magnitude in 2M.

**Fig. 6.**
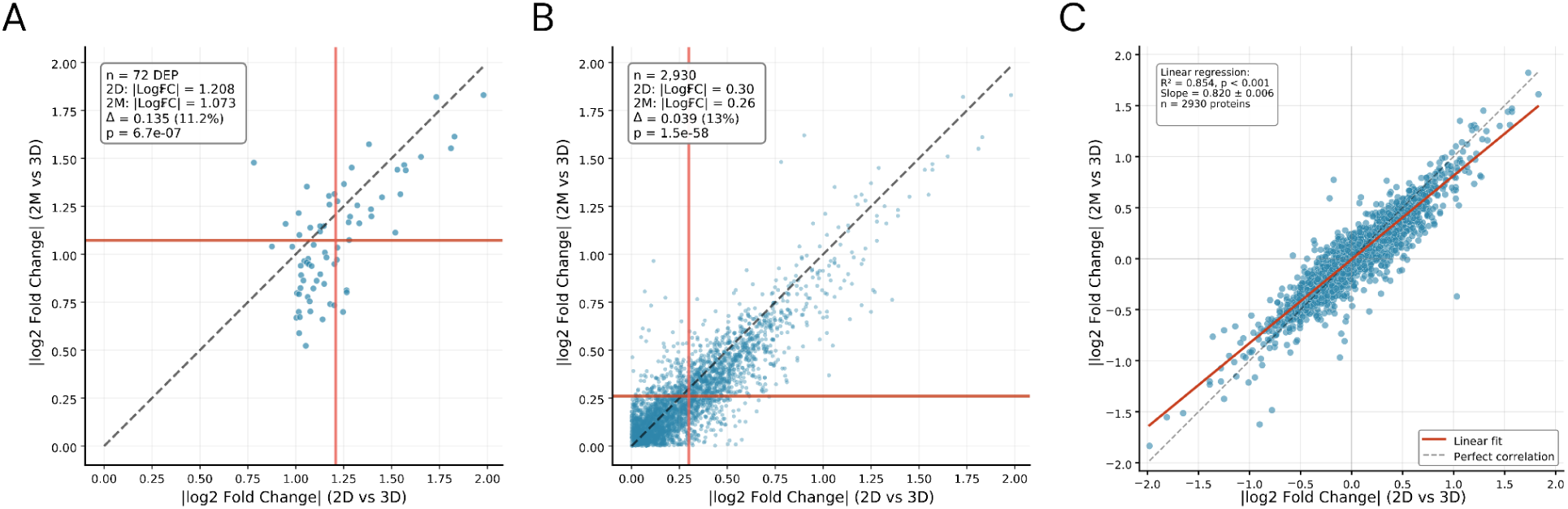
Matrigel contact alone induces a subtle but significant proteome-wide shift towards the 3D phenotype. **(A)** Comparison of the absolute Log₂(Fold Change) values for the 72 DEPs relative to the 3D condition. The magnitude of change is significantly smaller for the 2M vs. 3D comparison than for the 2D vs. 3D comparison (mean |ΔLog₂FC| = 0.135, paired t-test p = 6.7e-07), indicating the 2M proteome is closer to the 3D state. **(B)** The same analysis extended to all 2,930 non-missing proteins confirms the trend. The average fold-change magnitude is 13% smaller in the 2M condition (mean |ΔLog_₂_FC| = 0.039, paired t-test p = 1.5e-58). In both plots, the dashed line represents identity (x=y), and red lines indicate the mean values. **(C)** Linear regression of the signed Log₂(Fold Change) values for all 2,930 proteins. The slope of 0.82 (R² = 0.854, p < 0.001) demonstrates that while the direction of regulation is highly concordant, the magnitude of change in the 2M condition is systematically compressed by ∼18% compared to the 2D condition, relative to 3D.

A signed-regression of Log_₂_FC (2M vs 3D) on log2FC (2D vs 3D) gave a slope of 0.82 ± 0.006 (Fig. 6C), further indicating that cells merely contacting Matrigel exhibited on average a 18 % smaller proteomic shift towards the 3D phenotype when compared to cells on the 2D group . This trend, however, was not universal, as directionality was concordant for 2,579 of 2,930 proteins (88%).

While the statistical significance of this shift is profound, its effect size is modest. The large number of proteins in the paired test provides high statistical power, leading to a highly significant p-value even with a small underlying effect. The calculated Cohen’s d of 0.30 confirms this, indicating the effect is a weak, proteome-wide recalibration rather than a large-scale reprogramming. Taken together, these statistical and PCA results indicate that Matrigel interface alone does not trigger large-scale differential expression on the intermediate 2M group compared to a standard 2D monolayer, contact with Matrigel alone exerts some influence on the cell proteome, nudging it closer to the 3D group’s phenotype.

## DISCUSSION

The shift towards more physiologically relevant *in vitro* systems is crucial for advancing our understanding of disease and improving the predictiveness of preclinical drug screening. Three-dimensional cultures, which better preserve native epithelial cell states compared to traditional 2D monolayers, are at the forefront of this effort, providing more reliable models for precision medicine and reducing the reliance on animal testing. In this study, we provide a proteomic characterization of the widely used Calu-3 lung cell line, as it transitions from a classic two-dimensional (2D) monolayer to a three-dimensional (3D) spheroid architecture using Matrigel-GFR as the supporting scaffold, while also including an intermediate condition (2M) of monolayers cultured in the presence of Matrigel-GFR. We also executed a methodological framework capable of deconvolving the distinct contributions of physical dimensionality versus the biochemical cues inherent to the Matrigel scaffold. Our data revealed that while the 3D architecture is the primary driver of proteomic remodeling, the biochemical signals from the Matrigel-GFR substrate alone induced a subtle, yet statistically significant, systemic shift in the cellular proteome mostly towards the 3D phenotype.

A core strength of this work, and a point that warrants emphasis, lies in the careful consideration of some of the details in the experimental design, which was crafted to isolate variables and enhance physiological relevance. For instance, our use of a DMEM formulation with 1.8 g/L sodium bicarbonate, rather than the conventional 3.7 g/L, aligns with modern evidence-based guidelines for maintaining a stable physiological pH of ∼7.4 in incubators with 5% CO2 (41). This is not a trivial detail; it ensures that observed phenotypic changes are not confounded by pH stress, a common but often overlooked artifact in cell culture. Furthermore, the decision to set the culture period to seven days was deliberate. Unlike typical short-term 2D culture experiments (24-72 hours), this duration allowed the 3D spheroids to mature and establish stable, lumen-forming structures (42) while ensuring, through careful initial seeding density optimization, that the 2D and 2M monolayer cultures reached confluence only at the point of harvest. This synchronization is critical, as it minimizes quiescence-related artifacts and allows for a more synchronous comparison between biological states, as longitudinal studies have demonstrated that spheroid proteomes are dynamic and evolve over time, with very early spheroids (e.g., day 2) remaining proteomically similar to 2D monolayers (43).

One key methodological strength of our study, enabling the confident dissection of these proteomic shifts, was our rigorous two-level strategy for managing Matrigel contamination. At the experimental level, acknowledging that residual matrix proteins can influence quantitative proteomics, we first employed dispase as the reagent of choice for cell harvesting, the experimentally superior method for minimizing matrix carry-over, as reported by Wang et al. 2022 (13), who demonstrated its highest peptide yield and a SILAC incorporation ratio of 97.1%, indicating minimal leftover matrix contamination. In addition to the dispase method we also employed repeated washes of the samples with ice-cold DPBS solution to facilitate Matrigel liquefaction and removal. We complemented this strategy at the data analysis level, by searching our spectra against a combined human-mouse database and further removing known high-confidence Matrigel contaminants. This approach likely led to the exclusion of some legitimate human proteins that share high sequence homology with their mouse counterparts. However, this trade-off was a conscious, conservative choice, prioritizing the reduction of false-positive human-identifications and inflated ion intensity signal by Matrigel contaminants, over maximizing the sheer number of identified proteins. This combined strategy ensures that the final proteomic dataset generated is free of technical artifacts from matrix contamination, improving the confidence of downstream analysis.

The most striking observation was the extensive proteomic remodeling in 3D spheroids compared to both 2D and 2M monolayers even under our stringent filtering steps and cutoff thresholds. The functional enrichment analysis of proteins significantly altered by the 3D environment points to a shift from a proliferative, surface-exposed state to a more stable, tissue-like structure.

The significant downregulation in 3D of proteins associated with “Virus receptor activity,” “Cell surface,” and “Integral component of plasma membrane” is a classic indicator of this transition. Cells in a 2D monolayer are artificially polarized with a large apical surface area constantly interacting with the medium. In contrast, cells within a 3D spheroid establish extensive cell-cell junctions, reducing the exposed surface area and recapitulating a more native epithelial organization. This architectural change logically reduces the demand for certain surface receptors. Similarly, the downregulation of pathways involved in cholesterol and steroid biosynthesis likely reflects a shift in membrane biology away from the demands of rapid 2D proliferation towards a more homeostatic state within the spheroid.

Conversely, the “3d_up” protein set, which includes proteins exclusively found in the spheroids, was uniquely enriched for proteins involved with iron clusters. These cofactors are critical for a host of enzymes, particularly those in mitochondrial complexes involved in cellular respiration. This suggests a potential metabolic reprogramming in the 3D spheroids towards more efficient oxidative phosphorylation, a state often associated with more differentiated or quiescent cells compared to the highly glycolytic phenotype common in 2D cancer cultures. Furthermore, as some of these iron-sulfur proteins are involved in redox sensing and DNA repair, their upregulation may also reflect an adaptive response to micro-gradients of nutrients and hypoxia (44) that develop within the core of the spheroid structure.

Perhaps the most nuanced finding of this study is the quantitative evidence for a ’2M’ intermediate state that illustrates Matrigel’s non-spatial influence on cell phenotype. Our initial PCA clearly showed that 3D architecture is the dominant driver of proteomic change. However, it also revealed that the biochemical cues from Matrigel contact alone were not sufficient to create a distinct proteomic cluster, with the effect being smaller than the sample-to-sample biological variability within the monolayer conditions. The fact that this condition yielded no differentially enriched proteins when compared directly to 2D culture, yet was demonstrably closer to the 3D state in our paired statistical analysis, points to a key biological insight.

We conclude that the biochemical cues from the Matrigel-GFR substrate, absent the overarching influence of 3D architecture, do not trigger large-scale changes in protein expression. Instead, they appear to exert a subtle, systemic ’nudge’ on the proteome. This effect is powerful enough to be statistically significant at a global, proteome-wide level but is insufficient to reliably overcome inherent biological noise on a sample-by-sample basis, as visualized in the PCA.

Our results provide quantitative validation for the empirically held view that both dimensionality and matrix biochemistry govern cell phenotype. While the influence of growth factors and other components in animal-derived matrices is well-accepted, our data experimentally supports this statement, demonstrating that while dimensionality is the main variable that forces major reprogramming, the cellular proteomics is also modulated by substrate biochemistry.

The field remains heavily reliant on animal-derived extracellular matrices like Matrigel, which, despite being the gold standard, suffer from batch-to-batch variability and high cost. Current research is actively exploring several Matrigel-free avenues, which broadly include decellularized native ECM, tunable synthetic hydrogels (e.g., PEG, alginate), and gel-forming recombinant proteins (45). A central challenge in designing those alternatives is finding the precise balance of biochemical ligands and mechanical properties required. Our findings suggest that the biochemical cues from Matrigel alone, absent the 3D architecture and mechanical cues, provide a subtle “nudge” to the proteome. Pinpointing these effects is a critical step towards developing defined, synthetic alternatives that can replicate the benefits of ECM without the inherent drawbacks (6).

While this study provides key insights, certain inherent limitations should be considered when interpreting the results. Although our characterization provides a valuable baseline it is important to emphasize that our findings are based on a single batch of Matrigel-GFR for all replicates, and the batch-to-batch variability of animal-derived matrices like Matrigel is well documented (5, 46, 47). Also, our intermediate condition (2M) was based on the mostly indirect interaction of the cell monolayer with the components of the Matrigel droplets that were either soluble and able to diffuse into the media or that would naturally decompose during the duration of the experiment. Future work incorporating an intermediate condition where the cell is grown as a monolayer on top of a Matrigel-coated surface would be invaluable to help further quantify a more broad chemical effect of the matrix in spheroid/organoid culture. We also acknowledge that although growth-factor-reduced Matrigel was used, a potential “dosage effect” from a higher effective concentration of remaining biomolecules in the immersive 3D environment cannot be fully excluded (48). Also, our use of DDA proteomics has known limitations in sampling stochasticity, and our bulk analysis reflects an average proteome, potentially masking subpopulation heterogeneity that is expected within the spheroids.

Our use of a Conditionally-Quantified Protein (CQP) analysis, alongside the traditional DEP approach, allowed us to capture another dimension of proteomic remodeling. However, we acknowledge the methodological considerations inherent in this approach. The intensity filter we applied, while empirically justified and crucial for enhancing confidence, is a pragmatic choice. We reasoned that any protein whose levels falls below this threshold is, for functional purposes, effectively ’off’ or present at a level too low enough to be biologically suggestive of differential abundance in that context, an interpretation consistent with the Log₂ fold-changes >1 observed in our DEP analysis (49).

In conclusion, this study provides a high-resolution proteomic blueprint of Calu-3 cell adaptation to different microenvironments. We demonstrate that 3D architecture is ultimately the main driver of the profound reprogramming of the cellular proteome and, importantly, provide quantitative evidence that the biochemical composition of the EHS matrix induces a subtle, but measurable shift towards most of the final 3D phenotype. This works serves as a critical resource for researchers, providing a detailed experimental design rationale to help guide the decision of whether the added complexity, robustness and intrinsic limitations of a 3D Matrigel model are warranted for their specific experimental questions in the context of lung cell biology and drug discovery, and provides a dataset that can aid the design of the next generation of a defined, fully synthetic matrix alternative for lung cell biology and 3D culture.

## ACKNOWLEDGEMENTS

This research used facilities at the National Biosciences Laboratory (LNBio), the National Center for Research in Energy and Materials (CNPEM), a Social Organization supervised by the Brazilian Ministry of Science, Technology and Innovations (MCTI). The staff from the Mass Spectrometry Laboratory are acknowledged for assistance during experiments (proposal ID 20232410).

## DATA AVAILABILITY

The mass spectrometry proteomics data have been deposited to the ProteomeXchange Consortium via the PRIDE partner repository with the dataset identifier PXD066407. All the scripts utilized for sorting, filtering, data analysis and plotting are available on GitHub repository https://github.com/arielmmaia/c3d_proteomics_scripts.git

## SUPPLEMENTAL DATA

This article contains supplemental data.

## AUTHOR CONTRIBUTIONS

**A.M.M.**: Conceptualization, Methodology, Software, Formal analysis, Investigation, Writing – Original Draft, Writing – Review & Editing, Visualization, Project administration **C.V.B.**: Conceptualization, Writing – Review & Editing, Supervision, Project administration, Funding acquisition **P.M.**: Funding acquisition **L.A.B.**: Funding acquisition

## FUNDING AND ADDITIONAL INFORMATION

This work was supported by the National Institute of Science and Technology on Tuberculosis – INCT-TB (CNPq-FAPERGS-CAPES) [grant numbers: 421703-2017-2 and 17-1265-8]. C. V. Bizarro, L. A. Basso, and P. Machado are Research Career Awardees of the Brazilian National Council for Scientific and Technological Development (CNPq). This study was also financed in part by the Brazilian Federal Agency for Support and Evaluation of Graduate Education (CAPES), Finance Code 001.

## ABBREVIATIONS

BCA: (bicinchoninic acid)
CQP: (conditionally quantified protein)
DDA: (data dependent acquisition)
DEP: (differentially enriched protein)
DMEM: (Dulbecco’s modified Eagle’s medium)
DPBS: (Dulbecco’s phosphate-buffered saline)
DTT: (dithiothreitol)
ECM: (extracellular matrix)
FBS: (fetal bovine serum)
FDR: (false discovery rate)
GFR: (growth factor reduced)
IAA: (iodoacetamide)
LC-MS/MS: (liquid chromatography-tandem mass spectrometry)
Log_₂_FC: (Log_₂_ Fold-Change)
MBR: (match between runs)
MS: (mass spectrometry)
NAMs: (New Approach Methodologies)
PCA: (Principal Component Analysis)

## REFERENCES

1. Fitzpatrick, L. E., and McDevitt, T. C. (2015) Cell-derived matrices for tissue engineering and regenerative medicine applications. Biomater. Sci. 3, 12–24

2. Bhuker, S., Sinha, A. K., Arora, A., Tuli, H. S., Datta, S., Saini, A. K., Saini, R. V., and Ramniwas, S. (2025) Genes and proteins expression profile of 2D vs 3D cancer models: a comparative analysis for better tumor insights. Cytotechnology 77, 51

3. FDA Announces Plan to Phase Out Animal Testing Requirement for Monoclonal Antibodies and Other Drugs | FDA (2025)

4. FDA no longer needs to require animal tests before human drug trials | Science | AAAS

5. Hughes, C. S., Postovit, L. M., and Lajoie, G. A. (2010) Matrigel: a complex protein mixture required for optimal growth of cell culture. Proteomics 10, 1886–1890

6. Aisenbrey, E. A., and Murphy, W. L. (2020) Synthetic alternatives to Matrigel. Nat. Rev. Mater. 5, 539–551

7. Vukicevic, S., Kleinman, H. K., Luyten, F. P., Roberts, A. B., Roche, N. S., and Reddi, A. H. (1992) Identification of multiple active growth factors in basement membrane Matrigel suggests caution in interpretation of cellular activity related to extracellular matrix components. Exp. Cell Res. 202, 1–8

8. Patel, R., and Alahmad, A. J. (2016) Growth-factor reduced Matrigel source influences stem cell derived brain microvascular endothelial cell barrier properties. Fluids Barriers CNS 13, 6

9. Szklarczyk, D., Kirsch, R., Koutrouli, M., Nastou, K., Mehryary, F., Hachilif, R., Gable, A. L., Fang, T., Doncheva, N. T., Pyysalo, S., Bork, P., Jensen, L. J., and von Mering, C. (2023) The STRING database in 2023: protein-protein association networks and functional enrichment analyses for any sequenced genome of interest. Nucleic Acids Res. 51, D638–D646

10. Nonnis, S., Maffioli, E., Zanotti, L., Santagata, F., Negri, A., Viola, A., Elliman, S., and Tedeschi, G. (2016) Effect of fetal bovine serum in culture media on MS analysis of mesenchymal stromal cells secretome. EuPA Open Proteom. 10, 28–30

11. Shin, J., Rhim, J., Kwon, Y., Choi, S. Y., Shin, S., Ha, C.-W., and Lee, C. (2019) Comparative analysis of differentially secreted proteins in serum-free and serum-containing media by using BONCAT and pulsed SILAC. Sci. Rep. 9, 3096

12. Sanchez-Guzman, D., Boland, S., Brookes, O., Mc Cord, C., Lai Kuen, R., Sirri, V., Baeza Squiban, A., and Devineau, S. (2021) Long-term evolution of the epithelial cell secretome in preclinical 3D models of the human bronchial epithelium. Sci. Rep. 11, 6621

13. Wang, M., Yu, H., Zhang, T., Cao, L., Du, Y., Xie, Y., Ji, J., and Wu, J. (2022) In-Depth Comparison of Matrigel Dissolving Methods on Proteomic Profiling of Organoids. Mol. Cell. Proteomics 21, 100181

14. Varnavides, G., Madern, M., Anrather, D., Hartl, N., Reiter, W., and Hartl, M. (2022) In Search of a Universal Method: A Comparative Survey of Bottom-Up Proteomics Sample Preparation Methods. J. Proteome Res. 21, 2397–2411

15. Smith, P. K., Krohn, R. I., Hermanson, G. T., Mallia, A. K., Gartner, F. H., Provenzano, M. D., Fujimoto, E. K., Goeke, N. M., Olson, B. J., and Klenk, D. C. (1985) Measurement of protein using bicinchoninic acid. Anal. Biochem. 150, 76–85

16. Avelino, T. M., García-Arévalo, M., Torres, F. R., Goncalves Dias, M. M., Domingues, R. R., de Carvalho, M., Fonseca, M. de C., Rodrigues, V. K. T., Leme, A. F. P., and Figueira, A. C. M. (2022) Mass spectrometry-based proteomics of 3D cell culture: A useful tool to validate culture of spheroids and organoids. SLAS Discov. 27, 167–174

17. Rappsilber, J., Mann, M., and Ishihama, Y. (2007) Protocol for micro-purification, enrichment, pre-fractionation and storage of peptides for proteomics using StageTips. Nat. Protoc. 2, 1896–1906

18. Kovalchik, K. A., Colborne, S., Spencer, S. E., Sorensen, P. H., Chen, D. D. Y., Morin, G. B., and Hughes, C. S. (2019) Rawtools: rapid and dynamic interrogation of orbitrap data files for mass spectrometer system management. J. Proteome Res. 18, 700–708

19. Martens, L., Chambers, M., Sturm, M., Kessner, D., Levander, F., Shofstahl, J., Tang, W. H., Römpp, A., Neumann, S., Pizarro, A. D., Montecchi-Palazzi, L., Tasman, N., Coleman, M., Reisinger, F., Souda, P., Hermjakob, H., Binz, P.-A., and Deutsch, E. W. (2011) mzML--a community standard for mass spectrometry data. Mol. Cell. Proteomics 10, R110.000133

20. Chambers, M. C., Maclean, B., Burke, R., Amodei, D., Ruderman, D. L., Neumann, S., Gatto, L., Fischer, B., Pratt, B., Egertson, J., Hoff, K., Kessner, D., Tasman, N., Shulman, N., Frewen, B., Baker, T. A., Brusniak, M.-Y., Paulse, C., Creasy, D., Flashner, L., and Mallick, P. (2012) A cross-platform toolkit for mass spectrometry and proteomics. Nat. Biotechnol. 30, 918–920

21. Frankenfield, A. M., Ni, J., Ahmed, M., and Hao, L. (2022) Protein contaminants matter: building universal protein contaminant libraries for DDA and DIA proteomics. J. Proteome Res. 21, 2104–2113

22. Kong, A. T., Leprevost, F. V., Avtonomov, D. M., Mellacheruvu, D., and Nesvizhskii, A. I. (2017) MSFragger: ultrafast and comprehensive peptide identification in mass spectrometry-based proteomics. Nat. Methods 14, 513–520

23. Teo, G. C., Polasky, D. A., Yu, F., and Nesvizhskii, A. I. (2021) Fast deisotoping algorithm and its implementation in the msfragger search engine. J. Proteome Res. 20, 498–505

24. Yang, K. L., Yu, F., Teo, G. C., Li, K., Demichev, V., Ralser, M., and Nesvizhskii, A. I. (2023) MSBooster: improving peptide identification rates using deep learning-based features. Nat. Commun. 14, 4539

25. Käll, L., Canterbury, J. D., Weston, J., Noble, W. S., and MacCoss, M. J. (2007) Semi-supervised learning for peptide identification from shotgun proteomics datasets. Nat. Methods 4, 923–925

26. Demichev, V., Messner, C. B., Vernardis, S. I., Lilley, K. S., and Ralser, M. (2020) DIA-NN: neural networks and interference correction enable deep proteome coverage in high throughput. Nat. Methods 17, 41–44

27. da Veiga Leprevost, F., Haynes, S. E., Avtonomov, D. M., Chang, H.-Y., Shanmugam, A. K., Mellacheruvu, D., Kong, A. T., and Nesvizhskii, A. I. (2020) Philosopher: a versatile toolkit for shotgun proteomics data analysis. Nat. Methods 17, 869–870

28. Keller, A., Nesvizhskii, A. I., Kolker, E., and Aebersold, R. (2002) Empirical statistical model to estimate the accuracy of peptide identifications made by MS/MS and database search. Anal. Chem. 74, 5383–5392

29. Nesvizhskii, A. I., Keller, A., Kolker, E., and Aebersold, R. (2003) A Statistical Model for Identifying Proteins by Tandem Mass Spectrometry. Anal. Chem. 75, 4646–4658

30. Yu, F., Haynes, S. E., and Nesvizhskii, A. I. (2021) IonQuant Enables Accurate and Sensitive Label-Free Quantification With FDR-Controlled Match-Between-Runs. Mol. Cell. Proteomics 20, 100077

31. Ammar, C., Schessner, J. P., Willems, S., Michaelis, A. C., and Mann, M. (2023) Accurate Label-Free Quantification by directLFQ to Compare Unlimited Numbers of Proteomes. Mol. Cell. Proteomics 22, 100581

32. Tardif, M., Fremy, E., Hesse, A.-M., Burger, T., Couté, Y., and Wieczorek, S. (2023) Statistical Analysis of Quantitative Peptidomics and Peptide-Level Proteomics Data with Prostar. Methods Mol. Biol. 2426, 163–196

33. Kim, K.-Y., Kim, B.-J., and Yi, G.-S. (2004) Reuse of imputed data in microarray analysis increases imputation efficiency. BMC Bioinformatics 5, 160

34. Peng, H., Wang, H., Kong, W., Li, J., and Goh, W. W. B. (2024) Optimizing differential expression analysis for proteomics data via high-performing rules and ensemble inference. Nat. Commun. 15, 3922

35. Hsiao, Y., Zhang, H., Li, G. X., Deng, Y., Yu, F., Valipour Kahrood, H., Steele, J. R., Schittenhelm, R. B., and Nesvizhskii, A. I. (2024) Analysis and Visualization of Quantitative Proteomics Data Using FragPipe-Analyst. J. Proteome Res. 23, 4303–4315

36. Shah, A. D., Goode, R. J. A., Huang, C., Powell, D. R., and Schittenhelm, R. B. (2020) LFQ-Analyst: An Easy-To-Use Interactive Web Platform To Analyze and Visualize Label-Free Proteomics Data Preprocessed with MaxQuant. J. Proteome Res. 19, 204–211

37. Ritchie, M. E., Phipson, B., Wu, D., Hu, Y., Law, C. W., Shi, W., and Smyth, G. K. (2015) limma powers differential expression analyses for RNA-sequencing and microarray studies. Nucleic Acids Res. 43, e47

38. Benjamini, Y., and Hochberg, Y. (1995) Controlling the false discovery rate: a practical and powerful approach to multiple testing. Journal of the Royal Statistical Society: Series B (Methodological) 57, 289–300

39. Silva, J. M., Wippel, H. H., Santos, M. D. M., Verissimo, D. C. A., Santos, R. M., Nogueira, F. C. S., Passos, G. A. R., Sprengel, S. L., Borba, L. A. B., Carvalho, P. C., and Fischer, J. de S. da G. (2020) Proteomics pinpoints alterations in grade I meningiomas of male versus female patients. Sci. Rep. 10, 10335

40. Franciosa, G., Nieddu, V., Battistini, C., Caffarini, M., Lupia, M., Colombo, N., Fusco, N., Olsen, J. V., and Cavallaro, U. (2025) Quantitative proteomics and phosphoproteomics analysis of patient-derived ovarian cancer stem cells. Mol. Cell. Proteomics, 100965

41. Michl, J., Park, K. C., and Swietach, P. (2019) Evidence-based guidelines for controlling pH in mammalian live-cell culture systems. *Commun*. Biol. 2, 144

42. Qu, F., Zhao, S., Cheng, G., Rahman, H., Xiao, Q., Chan, R. W. Y., and Ho, Y.-P. (2021) Double emulsion-pretreated microwell culture for the in vitro production of multicellular spheroids and their in situ analysis. Microsyst. Nanoeng. 7, 38

43. Whitney, C. B., Beller, N. C., Fries, B. D., Lopez, A., and Hummon, A. B. (2025) Longitudinal proteomic changes in HCT 116 colon cancer spheroids during growth. J. Proteome Res.,

44. Lui, I., Schaefer, K., Kirkemo, L. L., Zhou, J., Perera, R. M., Leung, K. K., and Wells, J. A. (2025) Hypoxia induces extensive protein and proteolytic remodeling of the cell surface in pancreatic adenocarcinoma (PDAC) cell lines. J. Proteome Res.,

45. Kozlowski, M. T., Crook, C. J., and Ku, H. T. (2021) Towards organoid culture without Matrigel. *Commun*. Biol. 4, 1387

46. Hughes, C. S., Radan, L., Betts, D., Postovit, L. M., and Lajoie, G. A. (2011) Proteomic analysis of extracellular matrices used in stem cell culture. Proteomics 11, 3983–3991

47. Hughes, C., Radan, L., Chang, W. Y., Stanford, W. L., Betts, D. H., Postovit, L.-M., and Lajoie, G. A. (2012) Mass spectrometry-based proteomic analysis of the matrix microenvironment in pluripotent stem cell culture. Mol. Cell. Proteomics 11, 1924–1936

48. Sheth, M., Sharma, M., Lehn, M., Reza, H., Takebe, T., Takiar, V., Wise-Draper, T., and Esfandiari, L. (2024) Three-dimensional matrix stiffness modulates mechanosensitive and phenotypic alterations in oral squamous cell carcinoma spheroids. APL Bioengineering 8, 036106

49. Ghassami, A., Salehkaleybar, Kiyavash, and Bareinboim (2018) Budgeted Experiment Design for Causal Structure Learning. Bioinformatics 27,

