## Supplementary material for "From 2D Monolayer to 3D Spheroid: Matrigel’s Spatial and Chemical Cues Remodel the Calu-3 Proteome": Suplemental Data

### Table of contents

#### Supplemental Tables (excel spreadsheets)

**Supplemental Table S1** FragPipe-Analyst output with Limma analysis.

**Supplemental Table S2** Differentially Enriched Proteins (DEPs) in Calu-3 cells across culture conditions

**Supplemental Table S3** Conditionally-Quantified Proteins (CQPs) in Calu-3 cells across culture conditions

#### Supplemental Figures

**Supplemental Fig. S1** Waterfall plot showing the number of proteins identified across the nine samples

**Supplemental Fig. S2** Density plots of $Log_2$(Intensity) distributions for all quantified proteins within each condition (2D, 2M, 3D)

**Supplemental Fig. S3** Intra-condition reproducibility.

**Supplemental Fig. S4** PCA loadings plot for the analysis shown in Fig. 2C

**Supplemental Fig. S5** Rationale and filtering of Conditionally Quantified Proteins (CQPs).

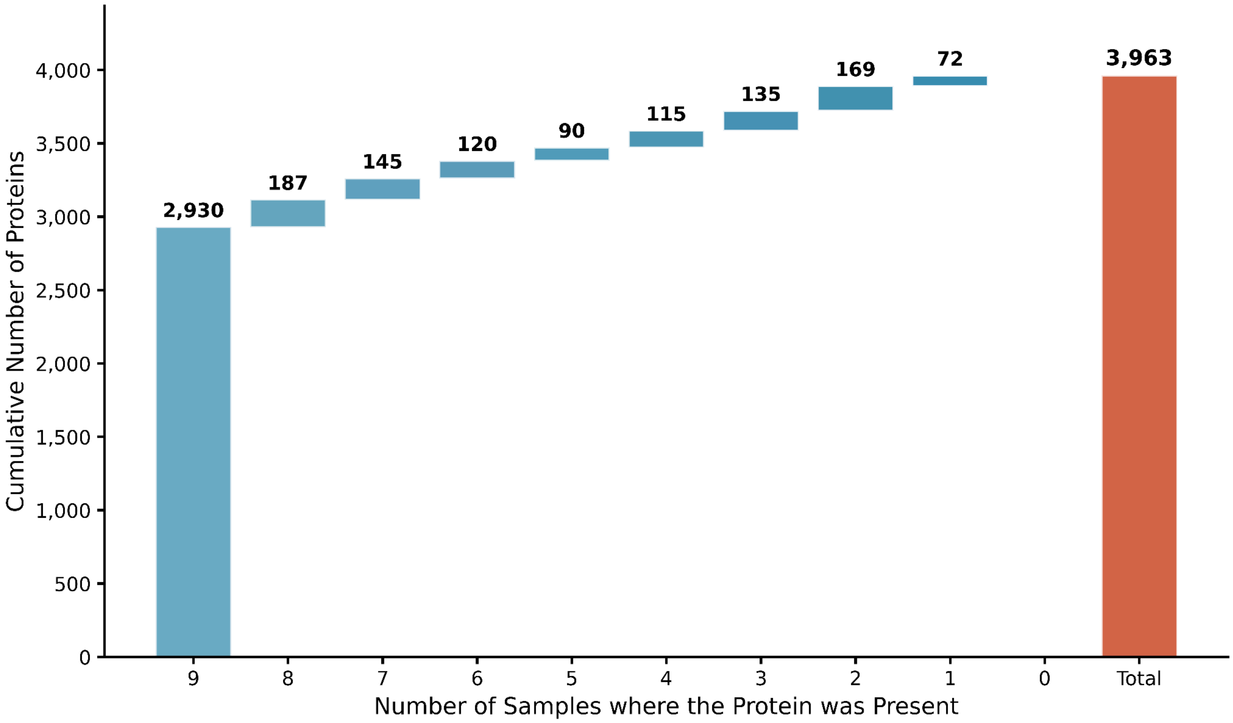


**Supplemental Fig. S1 Waterfall plot showing the number of proteins identified across the nine samples.** A core of 2,930 proteins was quantified in all nine samples, with a total of 3,963 proteins quantified in at least one sample.

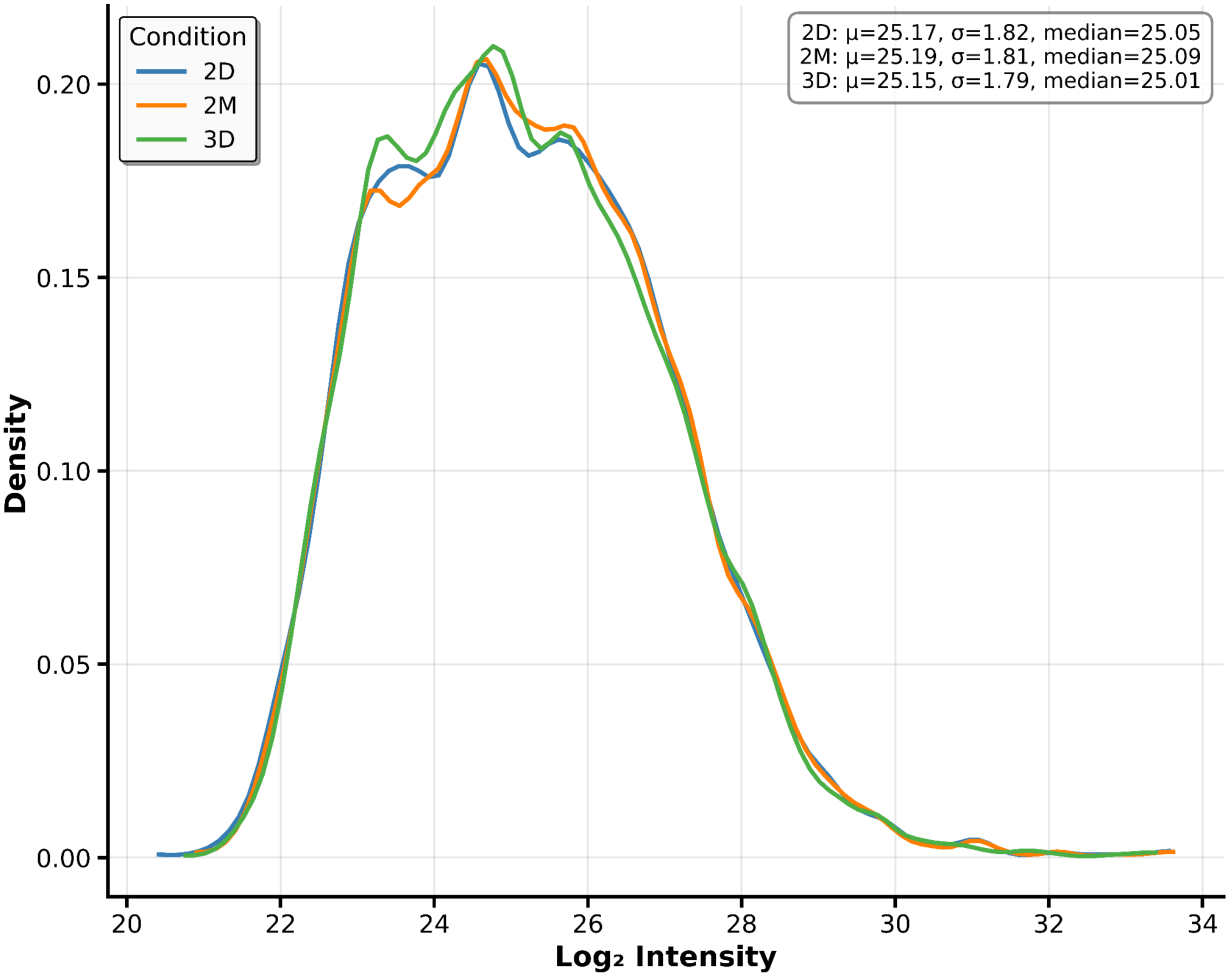


**Supplemental Fig. S2 Density plots of $Log_2$(Intensity) distributions for all quantified proteins within each condition (2D, 2M, 3D).** The highly overlapping distributions indicate comparable data quality and effective normalization across conditions.

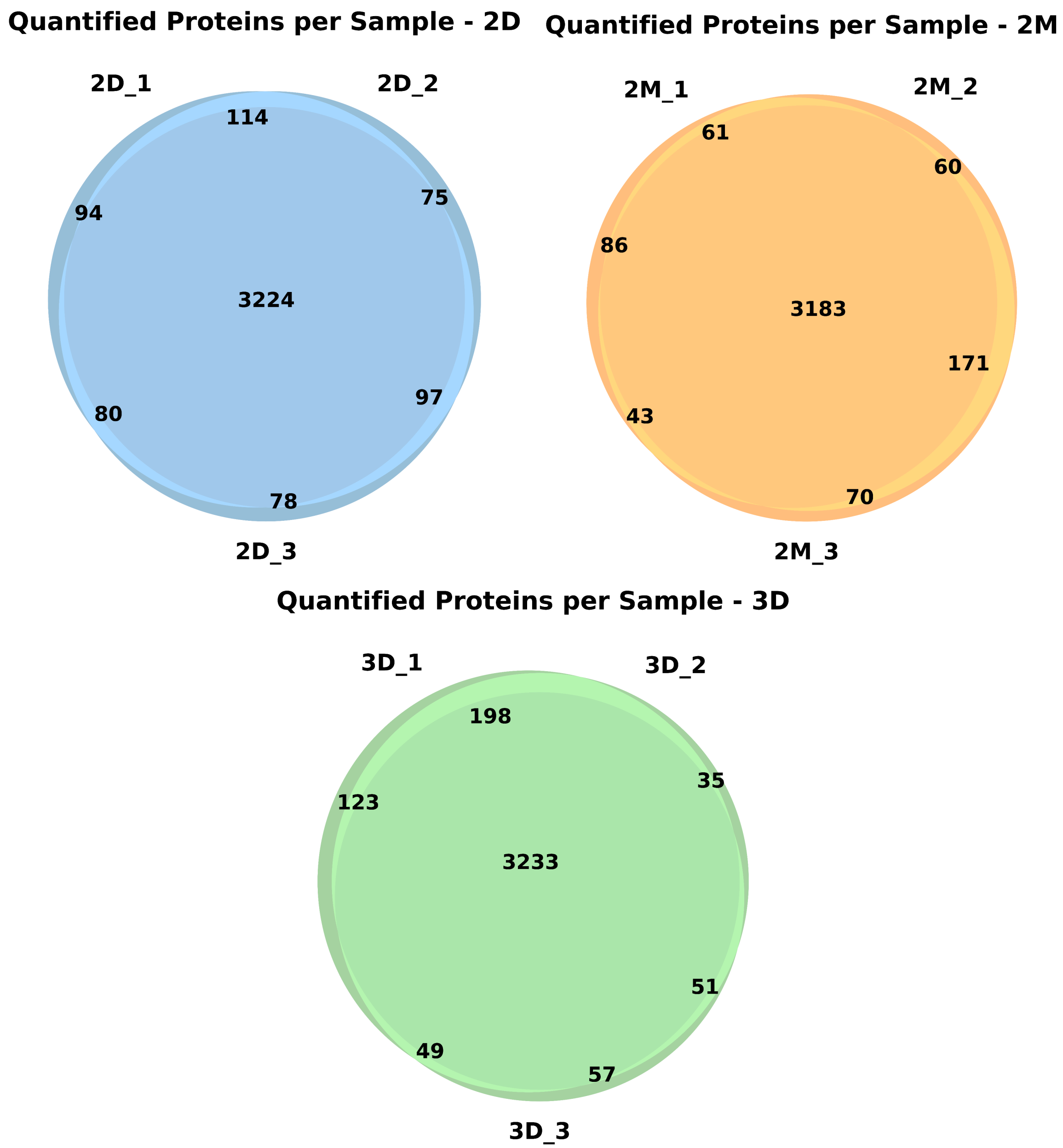


**Supplemental Fig. S3 Intra-condition reproducibility**. Venn diagrams showing the overlap of quantified proteins among the three biological replicates for each condition, demonstrating high intra-condition reproducibility.

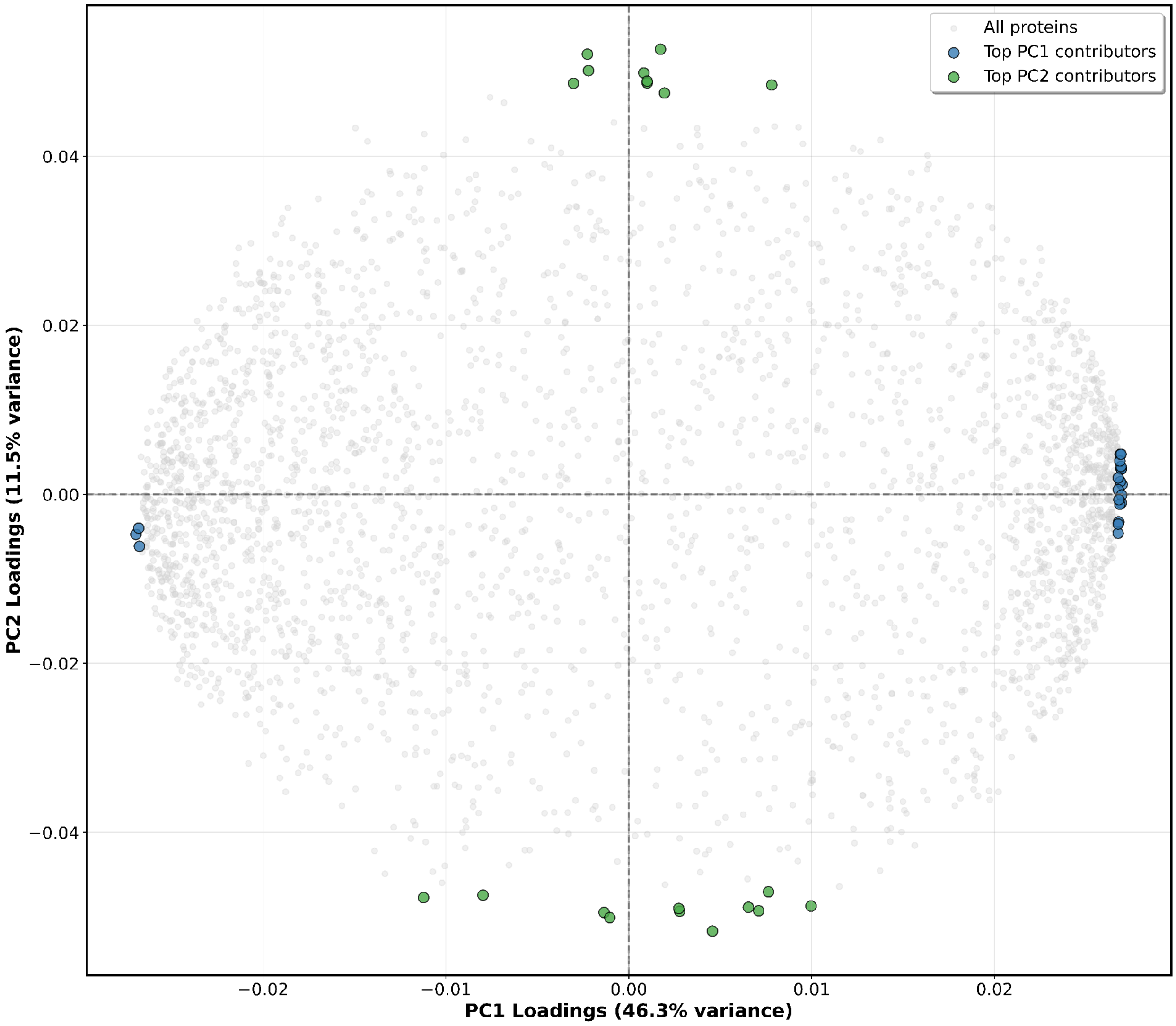


**Supplemental Fig. S4 PCA loadings plot for the analysis shown in Fig. 2C**. Proteins with the largest contribution to the variance explained by PC1 and PC2 are highlighted.

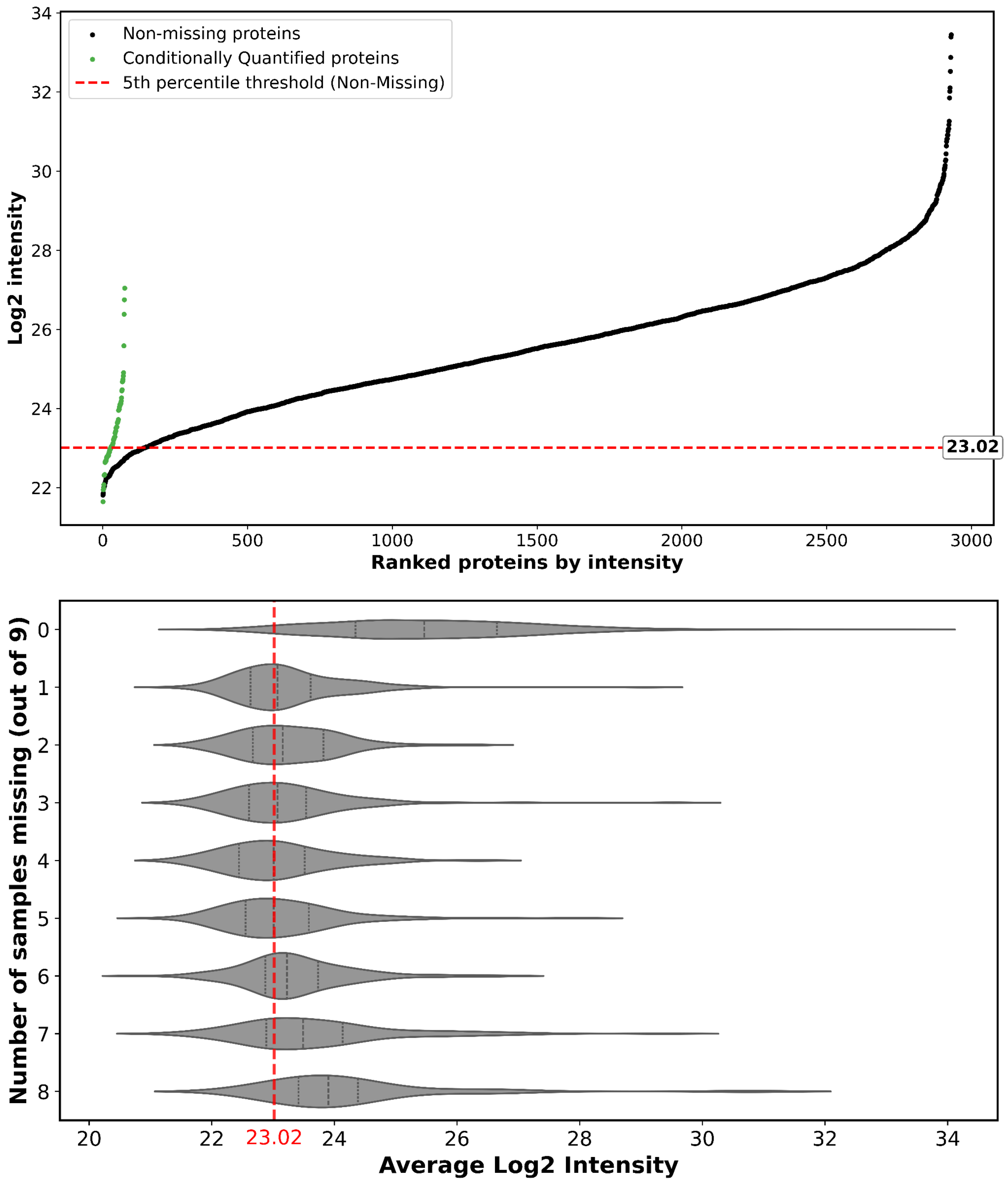


**Supplemental Fig. S5 Rationale and filtering of Conditionally Quantified Proteins (CQPs). (top)** Protein rank plot of average $Log_2$(Intensity) for the 2,930 non-missing proteins (black dots). The 76 CQPs are overlaid (green dots) to show their intensity distribution relative to the core proteome. **(bottom)** Violin plots illustrate the distribution of average $Log_2$(Intensity) for proteins stratified by the number of samples in which they were missing (out of 9 total samples). The plot demonstrates that proteins with lower average intensities are more frequently missing, a characteristic of shotgun proteomics data nearing the limit of detection. The red dashed line in both panels indicates the $Log_2$(Intensity) cutoff of 23.02 used to filter for high-confidence CQPs, corresponding to the 5th percentile of the non-missing protein dataset.
